# Generation of potent cellular and humoral immunity against SARS-CoV-2 antigens via conjugation to a polymeric glyco-adjuvant

**DOI:** 10.1101/2021.05.20.445060

**Authors:** Laura T. Gray, Michal M. Raczy, Priscilla S. Briquez, Tiffany M. Marchell, Aaron T. Alpar, Rachel P. Wallace, Lisa R. Volpatti, Maria Stella Sasso, Shijie Cao, Mindy Nguyen, Aslan Mansurov, Erica Budina, Elyse A. Watkins, Ani Solanki, Nikolaos Mitrousis, Joseph W. Reda, Shann S. Yu, Andrew C. Tremain, Ruyi Wang, Vlad Nicolaescu, Kevin Furlong, Steve Dvorkin, Balaji Manicassamy, Glenn Randall, D. Scott Wilson, Marcin Kwissa, Melody A. Swartz, Jeffrey A. Hubbell

**Affiliations:** Pritzker School of Molecular Engineering, University of Chicago; Chicago, IL 60637, United States; Committee on Immunology, University of Chicago; Chicago, IL 60637, United States; Animal Resources Center, University of Chicago; Chicago, IL 60637, United States; Department of Microbiology, Howard T. Ricketts Laboratory, University of Chicago; Chicago, IL 60637, United States; Department of Microbiology and Immunology, University of Iowa; Iowa City, IA 52242, United States; Department of Biomedical Engineering, Johns Hopkins School of Medicine; Baltimore, MD 21231, United States; Committee on Cancer Biology, University of Chicago; Chicago, IL 60637, United States; Ben May Department of Cancer Research, University of Chicago; Chicago, IL 60637, United States

**Author notes:** These authors contributed equally to this work (Laura T. Gray, Michal M. Raczy, Priscilla S. Briquez, Tiffany M. Marchell).

## Abstract

The SARS-CoV-2 virus has caused an unprecedented global crisis, and curtailing its spread requires an effective vaccine which elicits a diverse and robust immune response. We have previously shown that vaccines made of a polymeric glyco-adjuvant conjugated to an antigen were effective in triggering such a response in other disease models and hypothesized that the technology could be adapted to create an effective vaccine against SARS-CoV-2. The core of the vaccine platform is the copolymer p(Man-TLR7), composed of monomers with pendant mannose or a toll-like receptor 7 (TLR7) agonist. Thus, p(Man-TLR7) is designed to target relevant antigen-presenting cells (APCs) via mannose-binding receptors and then activate TLR7 upon endocytosis. The p(Man-TLR7) construct is amenable to conjugation to protein antigens such as the Spike protein of SARS-CoV-2, yielding Spike-p(Man-TLR7). Here, we demonstrate Spike-p(Man-TLR7) vaccination elicits robust antigen-specific cellular and humoral responses in mice. In adult and elderly wild-type mice, vaccination with Spike-p(Man-TLR7) generates high and long-lasting titers of anti-Spike IgGs, with neutralizing titers exceeding levels in convalescent human serum. Interestingly, adsorbing Spike-p(Man-TLR7) to the depot-forming adjuvant alum, amplified the broadly neutralizing humoral responses to levels matching those in mice vaccinated with formulations based off of clinically-approved adjuvants. Additionally, we observed an increase in germinal center B cells, antigen-specific antibody secreting cells, activated T follicular helper cells, and polyfunctional Th1-cytokine producing CD4^+^ and CD8^+^ T cells. We conclude that Spike-p(Man-TLR7) is an attractive, next-generation subunit vaccine candidate, capable of inducing durable and robust antibody and T cell responses.

## INTRODUCTION

Since December 2019, coronavirus disease 2019 (COVID-19), caused by severe acute respiratory syndrome coronavirus 2 (SARS-CoV-2), has evolved into a major global public health crisis. COVID-19 has overwhelmed global health systems due to its ease of transmission, considerable caseloads requiring hospitalization, long in-clinic recuperation times, and a confirmed case-mortality rate at around 1-5% (with significantly higher rates in patients with comorbidities and of older age) (*1–3*). As of May 2021, more than 150 million COVID-19 cases and more than 3 million deaths have been reported worldwide (*4*). These features highlight an urgent need for a vaccine against SARS-CoV-2.

SARS-CoV-2 infections begin through viral recognition of angiotensin-converting enzyme-2 (ACE2) on target cells (*5, 6*), mediated by the Spike glycoprotein that decorates the viral surface (*7, 8*). Spike typically exists as a homotrimer of 120 kDa proteins (>1100 residues each), of which the ACE2-binding function has been pinpointed to the receptor-binding domain (RBD) occurring around residues 319-541 (*9–11*). Therefore, interfering with this binding interaction, by generating antibodies against Spike and/or RBD, represents a promising strategy to limit viral infectivity (*12*), and in fact, has been the predominant approach used in today’s approved vaccines (*13–16*).

The urgent need for a vaccine has led to an immense number of vaccine candidates under various stages of development worldwide. As of May 2021, there were over 224 SARS-CoV-2 vaccine candidates under pre-clinical development and around 93 candidates in clinical trials (*17*). These numbers are the product of the inherent riskiness in the vaccine development process and include a wide range of technologies, such as DNA vaccines (*18*), vectored vaccines (*19, 20*), inactivated vaccines (*21*) and protein subunit vaccines (*22, 23*). Currently, two mRNA-loaded lipid nanoparticle formulations, developed by Pfizer-BioNTech (*24*) and Moderna (*25*), and one viral vector-based vaccine by Johnson&Johnson (*20, 26*) were granted emergency use authorizations in the US by the Food and Drug Administration in December 2020 and February 2021, respectively. Beyond the successes, there have also been notable disappointments in the race toward vaccine development, including Sanofi/GSK’s (*27*) and Merck’s (*28*) vaccine candidates that failed to elicit satisfactory immune responses in Phase 1/2 clinical trials. Given the continuing global pandemic, it is likely that more vaccine candidates will be explored and tested in continued efforts to control additional outbreaks, reduce hospitalization and mortality related to infection, reduce vaccine related adverse events, and address newly-emerging strains of SARS-CoV-2.

Ultimately, a successful vaccine against SARS-CoV-2 will provide protection from infection and effectively block the development of severe COVID-19. To do that, it must not only generate high neutralizing antibody titers that can prevent the virus from binding to host cells (*29, 30*), but it should also induce robust and durable T cell responses (*31, 32*). In fact, elevated T cell levels have been shown to be important in fighting SARS-CoV-2 infection in recovering patients, while reduced T cell numbers have been observed in patients who had severe disease (*33–35*). In addition, a vaccine candidate should also favor the production of T helper cell type 1 (Th1) over T helper cell type 2 (Th2) responses, as the latter have been associated with side effects including lung disease and vaccine-associated enhanced respiratory disease (*36, 37*). Conversely, Th1-biased immune responses have been shown to be associated with enhanced protection against viral infection (*38–40*). Finally, because COVID-19 is disproportionately lethal for elderly patients (age > 65 years), an ideal vaccine must be effective in this age group, even though many vaccine candidates have decreased efficacy within this demographic (*41*).

Addressing these requirements, our group recently described a modular vaccine platform that incorporates a random co-polymer of mannose and imidazoquinoline toll-like receptor 7 (TLR7) agonist monomers (p(Man-TLR7)) with an antigen on the same macromolecule. This platform leverages the dendritic cell (DC)-targeting properties of mannose-binding C-type lectins to efficiently co-deliver antigens and a potent polymeric adjuvant to these cells, eliciting broad lymphocyte-driven responses (*42*). The simplicity of our platform design allows the reversible conjugation of amine-containing antigens to the synthetic polymer p(Man-TLR7) in a manner such that the native antigen is released after reduction and self-immolation of the linker in response to intracellular signals. Following administration, the immunogenic conjugates are successfully taken up by DCs, resulting in antigen processing, cross-presentation, and activation. p(Man-TLR7) successfully adjuvanted ovalbumin and the malaria circumsporozoite protein (CSP), eliciting robust and high-quality humoral and cellular immune responses (*42*). Moreover, vaccination with CSP-p(Man-TLR7) generated neutralizing antibodies that inhibited the invasion of *P. falciparum* sporozoites into human hepatocytes *ex vivo* (*42*).

In this work, we hypothesized that the success of p(Man-TLR7) as a vaccine platform in other disease models would translate to SARS-CoV-2, resulting in robust neutralizing antibody responses and T cell responses against a conjugated viral antigen. To explore this, p(Man-TLR7) was conjugated to either the prefusion-stabilized Spike protein or its RBD. To place our preclinical work into broader context, we also evaluated our Spike-p(Man-TLR7) vaccine against benchmark formulations based on the most clinically advanced subunit vaccine adjuvants.

## RESULTS

### *In vitro* characterization of antigen-p(Man-TLR7) conjugates

We first produced both Spike and RBD antigens and verified their binding ability to the ACE2 receptor via surface plasmon resonance (SPR; **Fig. S1, A and B**). The dissociation constants (K_d_) were quantified at 11.6 nM and 19.5 nM, respectively, which corresponded with reported values of 2.9-14.7 nM (*11, 43*) and 4.7-44.2 nM (*8, 9*), respectively. These antigens were then conjugated to the p(Man-TLR7) construct, yielding two subunit vaccine candidates: RBD-p(Man-TLR7) and Spike-p(Man-TLR7) (**Fig. 1A; Fig. S2, A and B; Fig. S3A**). The conjugation of the p(Man-TLR7) polymer to antigen via covalent self-immolative linkage was confirmed via sodium dodecyl sulfate-polyacrylamide gel electrophoresis (SDS-PAGE), as indicated by an increase in molecular weight (**Fig. 1B and Fig. S3B**). Because of the nature of this conjugation, the protein’s surface accessible lysine residues are modified at random, which could interfere with the binding ability to ACE2. As the ACE2 binding site on Spike and RBD is an important epitope for generating neutralizing antibodies (*12*), steric hindrance of this site by the p(Man-TLR7) polymer could negatively affect the generation of neutralizing antibodies. Despite these concerns, RBD-p(Man-TLR7) and Spike-p(Man-TLR7) both retained ACE2-binding activity, although at half the levels of unmodified antigens (**Fig. 1C and Fig. S3C**). Lastly, we validated that antigen-p(Man-TLR7) conjugates activated murine bone marrow-derived dendritic cells (BMDCs) in a manner consistent with previous publications (*42*). Unlike unmodified antigens, both RBD-p(Man-TLR7) and Spike-p(Man-TLR7) stimulated BMDCs to secrete the immunostimulatory cytokines IL-12p70, IL-6, and TNFα (**Fig. 1D and Fig. S3D**). Overall, functional recombinant Spike and RBD were successfully expressed in-house and coupled onto p(Man-TLR7) to generate conjugates that had retained ACE2 binding activity, but superior DC stimulation compared to unmodified antigen.

**Fig. 1.**
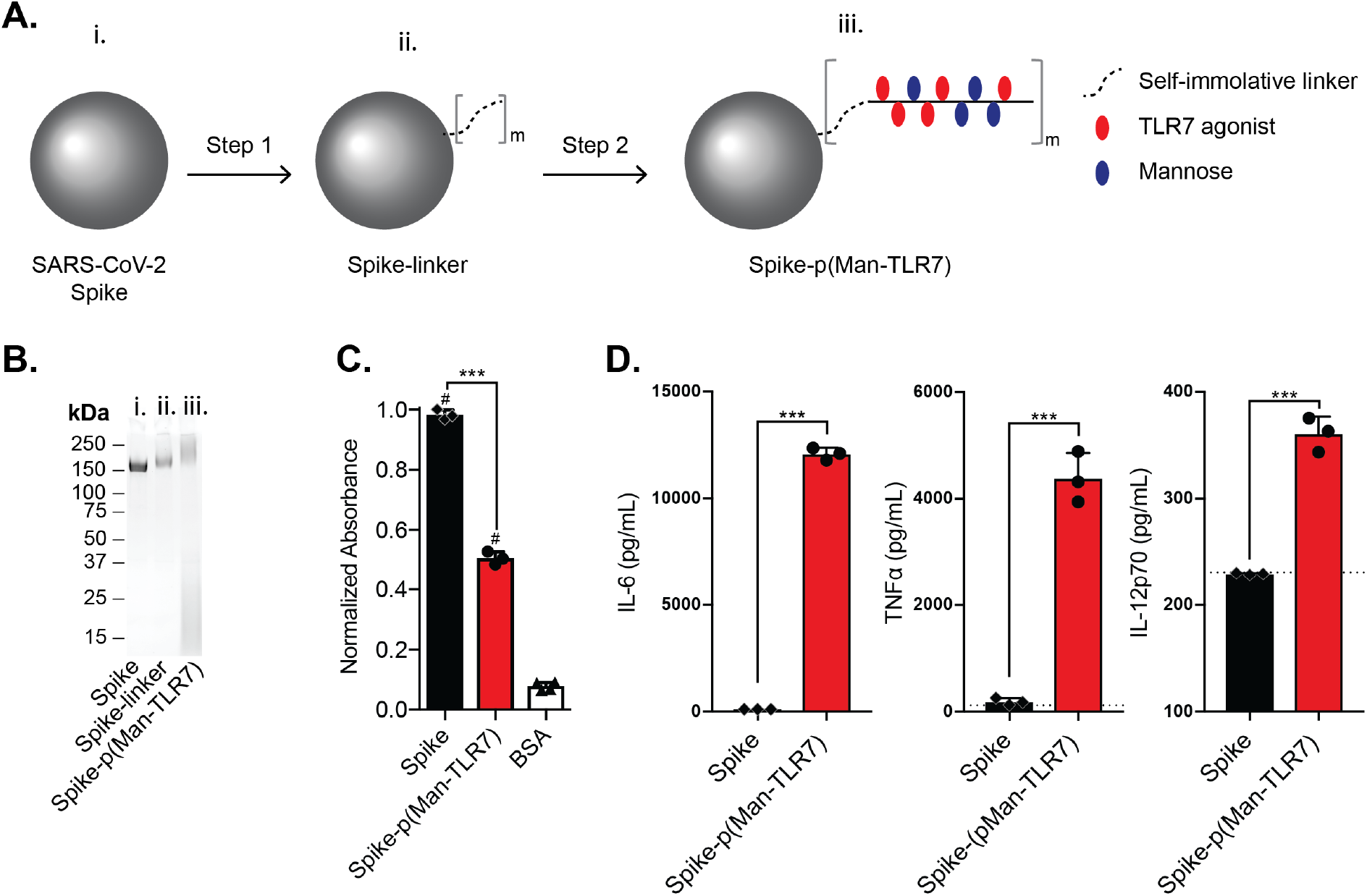
The prefusion-stabilized Spike antigen conjugated to p(Man-TLR7) is a potent activator of BMDCs. (**A**) Spike-p(Man-TLR7) is composed of Spike antigen (i.) conjugated, via a self-immolative linker (ii.), to a random copolymer synthesized from monomers that either activate TLR7 (red ovals) or target mannose-binding C-type lectins (blue ovals; iii.). (**B**) SDS-PAGE analysis of Spike before (i.) and after the two step conjugation reaction (ii., iii.). (**C**) Analysis of the binding ability of Spike-p(Man-TLR7) to human ACE2 (hACE2) via enzyme-linked immunosorbent assay (ELISA). (**D**) Concentration of IL-6, TNFα and IL-12p70 in the supernatant of BMDCs stimulated for 18h with either Spike or Spike-p(Man-TLR7) at the concentration corresponding to 25μM of the adjuvant, as determined by ELISA. Dotted horizontal lines represent the assay background. In (C and D), columns and error bars indicate mean+SD; statistical comparisons are based on one-way ANOVA with Tukey’s post-test: *** p<0.001; # p<0.001 as compared to bovine serum albumin (BSA).

### Vaccination with Spike-p(Man-TLR7) but not RBD-p(Man-TLR7) elicits SARS-CoV-2 neutralizing antibody responses

Next, we asked if the DC immune-stimulatory capacity of the conjugates *in vitro* would translate to superior antibody responses *in vivo*. To evaluate this, healthy adult C57BL/6 mice were vaccinated subcutaneously (s.c., in the hocks) in a prime-boost schedule 3 weeks apart, and sacrificed a week after the boost (**Fig. 2A and Fig. S4A**). We first assessed the humoral response in mice vaccinated with RBD-p(Man-TLR7) compared to mice vaccinated with RBD alone or adjuvanted with a mimic of the clinically-approved adjuvant AS04 (RBD+AS04-L; protein mixed with alum (aluminum hydroxide wet gel suspension) and monophosphoryl lipid A (MPLA); L for ‘like’), formulated according to published procedures (*44*). AS04 was designated as a positive control adjuvant due to its success in stimulating broad lymphocyte-driven responses against virally-mediated diseases such as the human papillomavirus (*45, 46*) and Hepatitis B virus (*47*), making it an ideal benchmark for evaluating the efficacy of the p(Man-TLR7) platform.

**Fig. 2.**
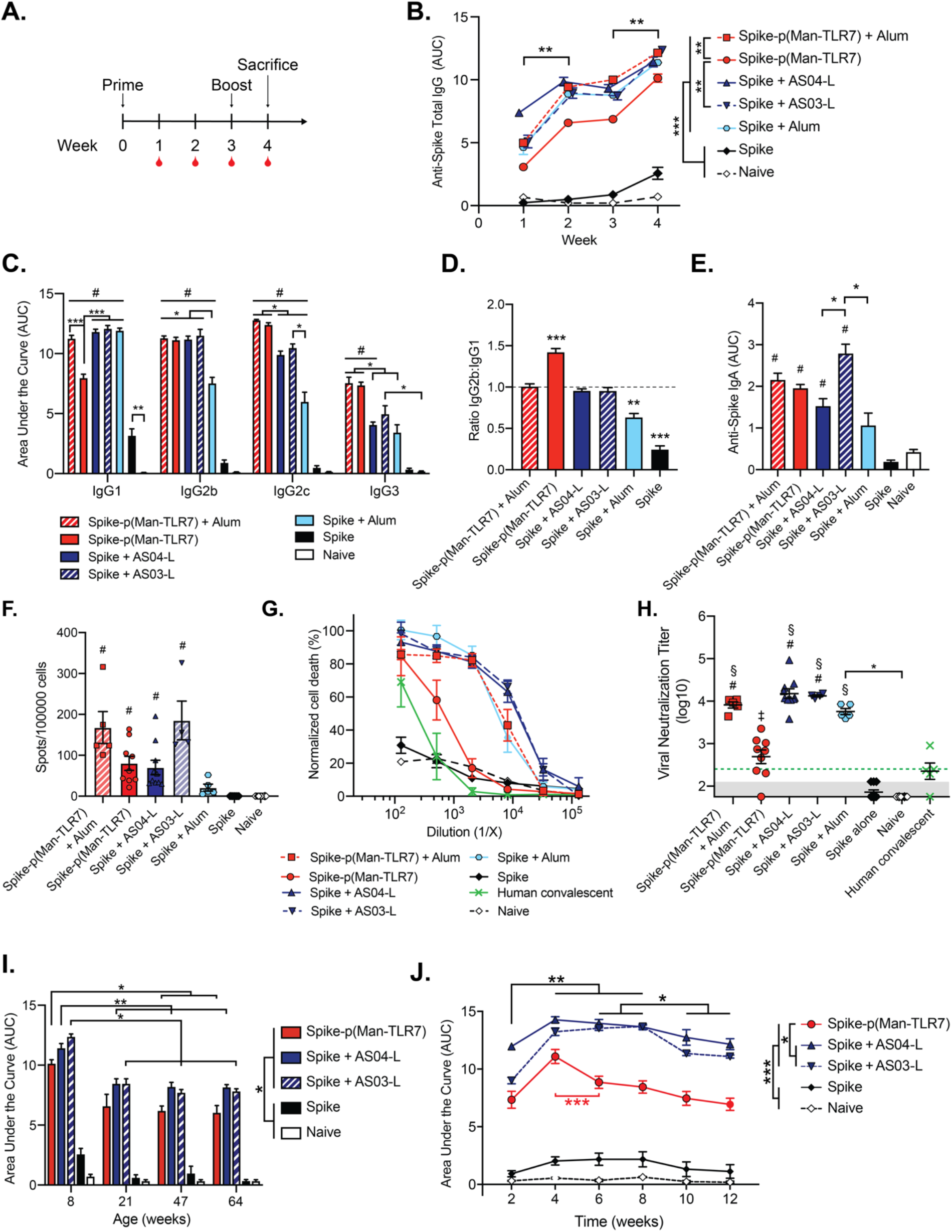
Spike-p(Man-TLR7) and Spike-p(Man-TLR7)+alum generate potent humoral responses in mice. (**A**) Mice were vaccinated with Spike-p(Man-TLR7), Spike-p(Man-TLR7)+alum, Spike+AS04-L, Spike+AS03-L, Spike+alum, or Spike at weeks 0 (prime) and 3 (boost), and their plasma was collected weekly until week 4. Plasma from naïve mice was collected at the same time points. (**B**) Total Spike-specific IgG antibodies over time reported as the area under the log-transformed curve (AUC) of absorbance vs. dilution. (**C**) Comparison of Spike-specific IgG isotypes (IgG1, IgG2b, IgG2c and IgG3) and (**D**) corresponding IgG2b:IgG1 ratios at one week post-boost (week 4). (**E**) Circulating anti-Spike IgA antibodies in the serum of vaccinated mice quantified at week 4 using AUC analysis. (**F**) Quantification of Spike-specific IgG^+^ antibody secreting cells by enzyme-linked immunosorbent spot (ELISpot) assay with splenocytes (Kruskal-Wallis with Dunn’s post-test). (**G**) Neutralization assay of SARS-CoV-2 infection on Vero-E6 cells in vitro. SARS-CoV-2 was pre-incubated with plasma isolated from mice at week 4. Percent neutralization was calculated based on viability of cells that did not receive virus (100%) or virus without plasma preincubation (0%). (**H**) Viral neutralization titers, representing plasma dilution at which 50% of SARS-CoV-2-mediated cell death is neutralized. Shaded area represents the lower limit of detection (titer of 2.11); green dotted horizontal line represents the FDA recommendation for “high titer” classification (= 2.40). (**I**) Comparison of total Spike-specific IgG antibodies in the plasma of 8, 21, 47 and >64 week old mice that received the indicated vaccines, following the same schedule as in (A). (**J**) Change in total Spike-specific IgG antibodies over time in plasma of mice (n = 5) that received the indicated vaccines, following the same vaccination schedule as in (A). All data are presented as mean SEM with n = 4-10 mice per group, unless stated otherwise. Comparisons were made using (B, I, and J) two-way ANOVA with Tukey’s multiple comparison test, (C and E) Brown-Forsythe ANOVA with Dunnet’s T3 test, (D) one sample t test (compared to the theoretical value of 1, representing an unbiased Th1/Th2 response), or (F and H) Kruskal-Wallis non-parametric test with Dunn’s post-test: * p<0.05, **p<0.01, *** p<0.001; # p<0.05 (for comparison to both Spike and naïve groups). Additional comparisons were made in (H) using Wilcoxon signed rank test: § p<0.05 and ‡ p=0.11 (as compared to the FDA “high titer” classification). In (B), comparisons noted on the graph are between the indicated timepoints for all groups except Spike and Naïve, while comparisons noted in the legend are between the indicated groups at week 4. In (I), comparisons indicated in the legend are true for mice at each age. In (J), comparisons noted on the graph in black are between the indicated timepoints for all groups except Spike and Naïve, and comparisons indicated in red are only for Spike-p(Man-TLR7). Comparisons indicated in the legend of (J) are true for every timepoint. In (B, I, and J), only relevant statistical comparisons are shown.

Vaccination with RBD-p(Man-TLR7) induced circulating levels of RBD-specific IgGs that trended higher than levels observed in mice vaccinated with RBD alone and naïve mice (p = 0.15 and p = 0.11, respectively; **Fig. S4, B and C**). Although RBD+AS04-L induced even higher levels of RBD-specific antibodies, RBD-p(Man-TLR7)-elicited antibody isotypes were suggestive of Th1-skewing (**Fig. S4, D and E**), as observed by comparing the ratio of IgG2b to IgG1 (*48, 49*), as well as increased levels of anti-RBD serum IgA (**Fig. S4, F and G**). IgG2 isotypes in mice are known to exhibit potent anti-viral activity (*50, 51*), and SARS-CoV-2-specific serum IgA antibodies have been shown to rapidly increase after the onset of COVID-19 and to have neutralization potential (*52, 53*).

We then asked if vaccine-elicited RBD-specific antibodies could effectively neutralize SARS-CoV-2 virions, preventing their ability to infect Vero-E6 cells *in vitro*. We observed that while plasma from mice vaccinated with RBD-p(Man-TLR7) showed an increase in virus neutralization titer (VNT) compared to mice vaccinated with RBD alone, it failed to meet the FDA-recommended VNT threshold for COVID-19 convalescent plasma therapy (**Fig. S4, H and I**) (*54*). At the same time, we observed that plasma from mice vaccinated with the RBD+AS04-L formulation protected Vero-E6 cells against viral infection *in vitro* (**Fig. S4, H and I**).

Next, we assessed the humoral responses of mice vaccinated with Spike-p(Man-TLR7). We also compared this formulation against additional benchmarks mimicking clinically-approved vaccine formulations based on alum: Spike+alum and Spike-p(Man-TLR7)+alum. Alum has been shown to enhance antigen availability, activation of antigen presenting cells (APCs), and uptake by immune cells through the formation of a depot at the injection site (*55–58*). Additionally, alum is commonly used in combination with other adjuvants with direct immunostimulatory activity, as embodied by one COVID-19 vaccine candidate in clinical testing that formulates Spike with alum and the TLR9 agonist CpG (*59*). Based on these properties of alum, we hypothesized that it could synergize with Spike-p(Man-TLR7) to produce a strong humoral response. In addition, we compared our formulations with Spike alone and Spike+AS04-L, as well as with Spike+AS03-L (Spike mixed with an oil-in-water emulsion of α-tocopherol, squalene, and polysorbate 80). AS03-L is an analog of the clinical AS03 adjuvant, which has been investigated in clinical trials as a COVID-19 vaccine with the Spike protein as the antigen (*60*).

In our studies, vaccination with Spike-p(Man-TLR7) elicited higher titers of Spike-specific antibodies versus vaccination with Spike alone (p<0.001) or in naïve mice (p<0.001, **Fig. 2B and Fig. S5A**). The benchmark vaccine formulations Spike+AS03-L and Spike+AS04-L elicited even higher Spike-specific IgG titers, but these levels were matched by Spike-p(Man-TLR7)+alum (**Fig. 2B and Fig. S5A**). Compared to all of these groups, however, Spike-p(Man-TLR7)-elicited IgG isotypes were more suggestive of Th1 activity, based on the ratio of IgG2b to IgG1 (**Fig. 2, C and D**). Notably, this vaccine, with or without alum, also increased levels of Spike-specific serum IgA, as compared to mice vaccinated with Spike alone, Spike adjuvanted with alum or AS04-L, or naïve mice (**Fig. 2E and Fig. S5B**). However, vaccination with Spike+AS03-L stimulated the highest levels of serum IgA among all groups (**Fig. 2E and Fig. S5B**). In agreement with these Spike-specific antibody responses, all adjuvanted formulations also resulted in an increase in the number of Spike-specific antibody secreting cells (ASCs) as compared to mice vaccinated with Spike alone or naïve mice, with the highest numbers of ASCs observed in mice vaccinated with Spike-p(Man-TLR7)+alum and Spike+AS03-L (**Fig. 2F and Fig. S5C**).

All adjuvanted vaccine formulations led to demonstrable neutralization of SARS-CoV-2 infection on Vero-E6 cells. In this regard, plasma from mice vaccinated with Spike-p(Man-TLR7) exceeded the FDA-recommended VNT threshold for convalescent plasma therapy, demonstrating superior neutralization activity over human convalescent plasma and 1.6-fold greater neutralization activity versus plasma from mice vaccinated with Spike alone or from naïve mice (**Fig. 2, G and H**). While vaccination with either Spike+AS04-L, Spike+AS03-L, or Spike+alum all led to even greater VNTs (**Fig. 2, G and H**), co-formulation of Spike-p(Man-TLR7) with alum allowed this platform to match the VNTs elicited by these positive control benchmarks (**Fig. 2, G and H**).

We then asked if the efficacy of Spike-p(Man-TLR7) in eliciting strong humoral responses in adult healthy mice would also translate to elderly mice. Four weeks after they received the priming dose, mice of all ages, ranging from 8 to >64 weeks, exhibited high titers of anti-Spike IgGs in all adjuvanted groups assessed (**Fig. 2I**). Furthermore, these responses were durable in adult mice, persisting for at least 12 weeks after the priming dose, with area under the curve (AUC) values from log-transformed enzyme-linked immunosorbent assay (ELISA) absorbance plots for total anti-Spike IgG exceeding 5.0 at week 12 (**Fig. 2J**). The AUC levels peaked at above 10 by week 4, and faded slightly between weeks 4 and 6, but remained at similar levels up to week 12 (**Fig. 2J**).

Taken together, our observations indicate that Spike-p(Man-TLR7) is able to induce a robust humoral response in both adult and elderly mice and that antibodies induced by the p(Man-TLR7) platform persist for at least 12 weeks after the priming dose. Additionally, the humoral response is further increased by the addition of alum to Spike-p(Man-TLR7).

### Expansion of epitopic coverage upon Spike-p(Man-TLR7) vaccination

Viruses tend to mutate to evade even the most effective neutralizing antibodies, and as such, a vaccination strategy that can elicit broad epitope coverage might be important to control a mutable virus. We characterized the repertoires of Spike-specific antibodies raised by each vaccine formulation via peptide arrays of linear Spike epitopes. The peptide arrays were constructed based on the full linear amino acid sequence of the SARS-CoV-2 Spike protein (NCBI GenBank accession # QHD43416.1), encompassing 254 unique 15-mer overlapping peptides with 5-amino acid offsets.

While vaccination with unadjuvanted Spike resulted in antibodies that recognized only a limited number of epitopes, p(Man-TLR7) conjugation expanded the epitope coverage to linear epitopes corresponding to the ACE2 binding site of RBD (*61*) and two previously reported linear Spike epitopes shown to elicit neutralizing antibodies in COVID-19 patients (**Fig. 3**) (*62, 63*). Notably, the addition of alum to Spike-p(Man-TLR7) further expanded the breadth of recognized epitopes, matching or surpassing the breadth of epitopes recognized by antibodies from mice vaccinated with Spike+AS04-L, Spike+AS03-L, or Spike+alum (**Fig. 3**).

**Fig. 3.**
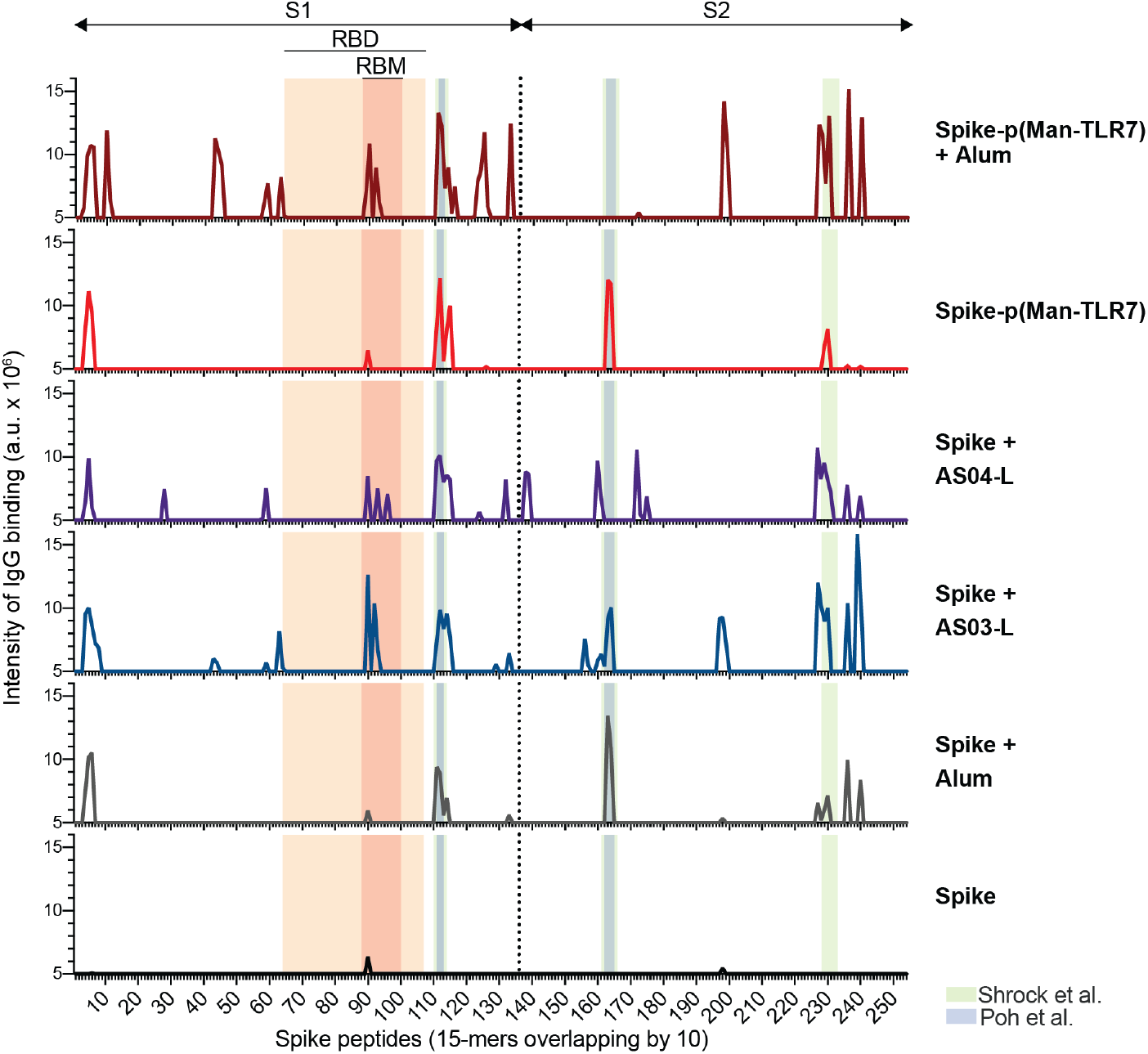
Vaccination with Spike-p(Man-TLR7) and Spike-p(Man-TLR7)+alum elicits a broad humoral response targeting the receptor binding motif (RBM) of RBD and other neutralizing linear epitopes. Mice were vaccinated as in Fig. 2A. Plasma was collected at week 4, pooled by vaccination group, and analyzed for binding to linear epitopes using a peptide array. X-axis represents the sequential peptide number within the Spike amino acid sequence (overlapping 15-amino acid peptides with 5-amino acid offsets). Y-axis quantifies the level of antibody binding to each peptide, detected via luminescence (a.u.). Axis begins from the value of the background, which was set at 5 × 10^6^ a.u. Several relevant regions of the Spike protein are indicated above the graphs: S1 and S2 subunits, RBD (light orange box), and RBM (dark orange box). Regions corresponding to neutralizing Spike epitopes identified by Shrock et al. *(62)* and Poh et al. *(63)* are indicated in light green and light blue boxes, respectively.

### Spike-p(Man-TLR7) platforms induce antigen-specific B cell immunity and expansion of T_fh_ cells

Due to the higher neutralizing antibody titers and broader epitope coverage found in the p(Man-TLR7) conjugated group compared to vaccination with Spike alone, we asked how these differences might be reflected in B cell responses in the secondary lymphoid organs. We analyzed the lymph nodes and spleens of vaccinated mice to examine the phenotypes and activation of B cells (**Fig. S6A**) and follicular helper CD4^+^ T (T_fh_) cells (**Fig. S8A**), the cells responsible for establishing humoral immunity.

Spike-p(Man-TLR7), both with and without alum, triggered B cell (CD19^+^ B220^+^) expansion in the draining lymph nodes and spleen as compared to mice vaccinated with Spike alone or naïve mice (**Fig. 4, A and B**). While Spike-p(Man-TLR7) elicited higher frequencies of germinal center (GC) B cells (IgD^−^ GL7^+^ CD38^−^) among splenic and lymph node B cells versus vaccination with Spike alone or in naïve mice, vaccines containing alum were generally even more effective at inducing GC B cells in the lymph nodes. As such, Spike-p(Man-TLR7)+alum triggered fourfold higher frequencies of GC B cells among lymph node B cells versus Spike-p(Man-TLR7) alone (3.4 ± 0.1% vs. 0.8 ± 0.1%) – levels matched by Spike+AS04-L (3.2 ± 0.5%) and Spike+alum (3.7 ± 0.4%) and exceeded by Spike+AS03-L (7.8 ± 0.5%; **Fig. 4, A and B**). In contrast, Spike-p(Man-TLR7) and Spike+AS03-L vaccination resulted in the highest frequencies of GC B cells in the spleen (1.2± 0.3% and 1.3± 0.3%, respectively), while the alum-containing formulations (Spike-p(Man-TLR7)+alum, Spike+AS04-L, and Spike+alum) did not result in as high levels of systemic GC responses (**Fig. 4B**). Reflective of these trends, the frequency of GC B cells that recognized RBD were elevated in mice treated with Spike-p(Man-TLR7) or Spike-p(Man-TLR7)+alum in the spleen or lymph nodes, respectively (**Fig. 4, A and B; Fig. S6, B and C**). Additionally, Spike-p(Man-TLR7)+alum increased the frequency of memory B cells (IgD^−^ GL7^−^ CD38^+^) in the draining lymph nodes compared to non-adjuvanted controls and to Spike-p(Man-TLR7) (**Fig. S7A**). Moreover, we observed a significant reduction in the naïve B cell population in the draining lymph nodes of mice vaccinated with either Spike-p(Man-TLR7) or Spike-p(Man-TLR7)+alum (**Fig. S7A**). In the spleen, the different vaccine formulations were not associated with statistically significant differences in memory or naïve B cell composition (**Fig. S7B**). Lastly, neither Spike-p(Man-TLR7) nor Spike-p(Man-TLR7)+alum increased the frequencies of plasmablasts (CD138^+^ B220^+^) or plasma cells (CD138^+^ B220^−^) via flow cytometry at one week post-boost in both the draining lymph nodes and spleen, although a trend towards increased levels was observed in the spleen (**Fig. S7, A and B**).

**Fig. 4.**
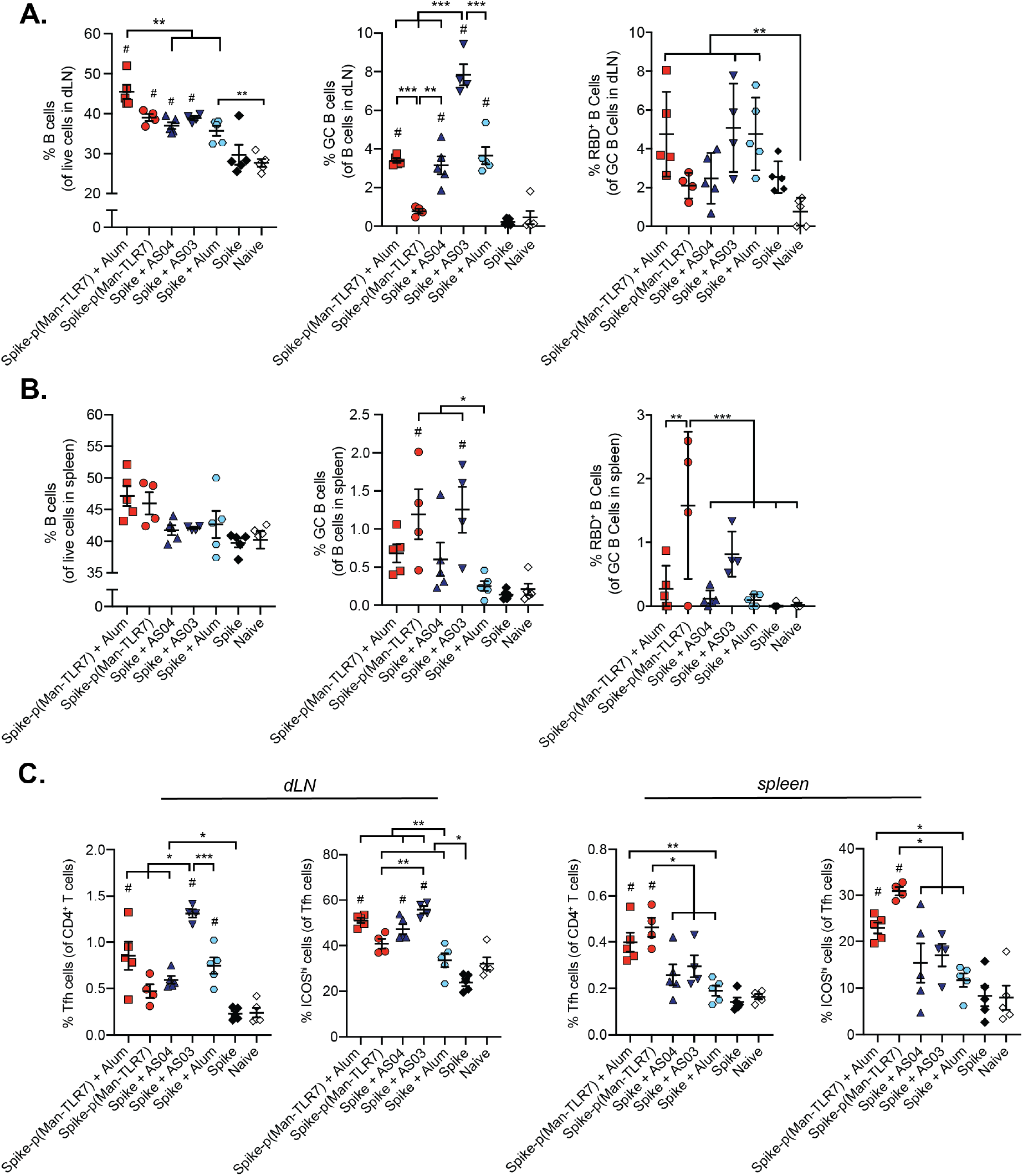
Secondary lymphoid organ-resident B cells and CD4^+^ T follicular helper (T_fh_) cells are activated in mice vaccinated with Spike-p(Man-TLR7) or Spike-p(Man-TLR7)+alum. (**A and B**) Quantification of B cells resident within the (A) draining LNs (dLNs) of the vaccination site or (B) spleen via flow cytometry at week 4 after vaccination as in Fig. 2A. From left to right, total B cells (CD19^+^ B220^+^) within live cells, germinal center (GC) B cells (IgD^−^GL7^+^CD38^−^) within live B cells, and RBD tetramer reactive GC B cells (RBD^+^) as a percentage of GC B cells. (**C**) Activation of CD4^+^ T_fh_ cells in the dLNs and the spleen was characterized by flow cytometry. T_fh_ cells were defined as CXCR5^+^BCL6^+^CD4^+^ and quantified within CD4^+^ T cells. ICOS^hi^ T_fh_ cells were quantified within T_fh_ cells. Data plotted as mean SEM with n = 4-5 mice per group. *p<0.05, ** p<0.01, ***p<0.001 by one-way ANOVA with Tukey’s post-test; # p<0.05 as compared to both Spike and naïve.

Shifting our focus to the T_fh_ cells (CD4^+^ Bcl6^+^ CXCR5^+^), we observed that animals vaccinated with Spike-p(Man-TLR7)+alum showed an increase in the fraction of T_fh_ cells in both the spleen and draining lymph nodes (dLNs) compared to mice treated with most other formulations (**Fig. 4C and Fig. S8A**). In addition, while a significantly higher fraction of these T_fh_ cells expressed a marker of activation (ICOS^hi^) in both the lymph nodes and spleen compared to unadjuvanted controls, we detected only modest trends towards increased proliferation of these cells, discerned by high expression of Ki67 (**Fig. 4C and Fig. S8B**). In the absence of alum, Spike-p(Man-TLR7) increased frequencies of both total and activated T_fh_ cells in the spleen, but not in dLNs, compared to mice treated with other formulations (**Fig. 4C and Fig. S8B**).

Altogether, Spike-p(Man-TLR7) vaccination induced antigen-specific B cell immunity and expansion of activated T_fh_ cells in the spleen, while the addition of the adjuvant alum localized the response to the dLNs.

### Th1 biased cellular responses are observed upon vaccination with Spike-p(Man-TLR7) with and without alum

The establishment of T cell responses plays an essential role in protection against infectious diseases (*64*). Some reports indicate that cellular immunity is as crucial as humoral immunity in COVID-19 recovery (*65*). Therefore, we characterized the antigen-specific T cell responses in the spleens of mice vaccinated with either Spike-p(Man-TLR7) conjugates or benchmark formulations. One week after the boost, splenocytes from all vaccinated and control mice were restimulated *ex vivo* with Spike peptide pools, and we quantified intracellular levels of the costimulatory cytokines IFNγ, TNFα, and IL-2 (**Fig. S9A**).

Vaccination with Spike-p(Man-TLR7) either alone or in combination with alum generated higher frequencies of cytokine^+^ CD4^+^ T cells, more polyfunctional CD4^+^ T cells (producing all three cytokines: IFNγ, TNFα, and IL-2), and higher expression of IFNγ compared to other groups (**Fig. 5, A and B; Fig. S9B**). Splenic CD8^+^ T cells elicited by Spike-p(Man-TLR7) vaccination trended towards increased intracellular cytokine expression relative to mice treated with Spike alone, but did not reach statistical significance (**Fig. 5, C and D; Fig. S9B**). Spike-p(Man-TLR7)+alum, however, resulted in a superior increase in cytokine^+^ CD8^+^ T cells and polyfunctional CD8^+^ T cells upon restimulation compared to other groups (**Fig. 5, C and D; Fig. S9B**).

**Fig. 5.**
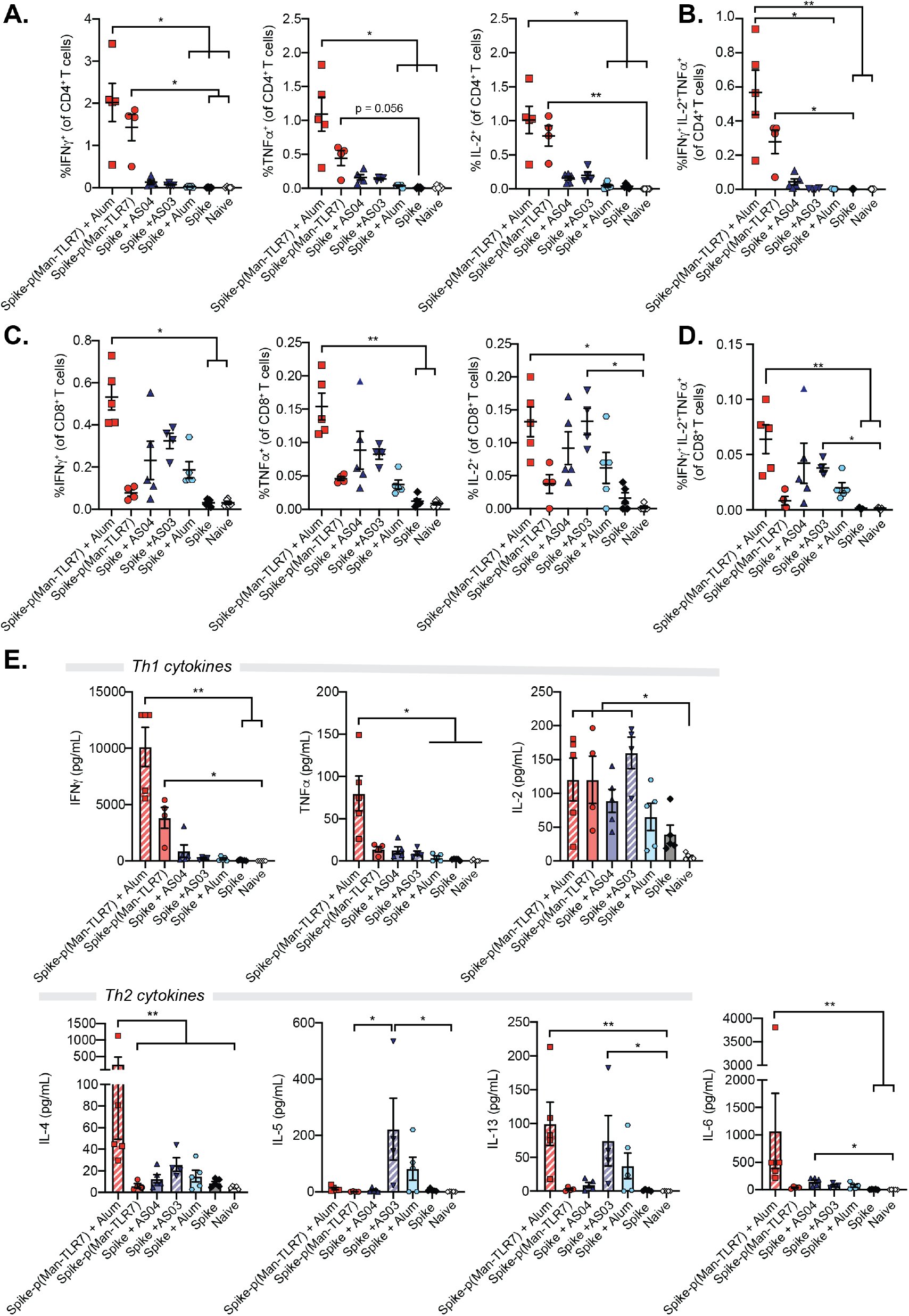
Vaccination with Spike-p(Man-TLR7) and Spike-p(Man-TLR7)+alum elicits robust antigen-specific T cells responses. Splenocytes harvested one week post-boost from vaccinated mice (vaccinated as in Fig. 2A) were restimulated ex vivo with a Spike-derived peptide pool for 6h prior to flow cytometry analysis or with the full length Spike protein for 3 days prior to multiplexed cytokine analysis. (**A to D**) Cytokine-producing (A and B) CD4^+^ and (C and D) CD8^+^ T cell responses were detected by intracellular staining and quantified by flow cytometry. (A) IFNγ +, TNFα^+^ and IL2^+^ CD4^+^ T cells quantified within CD4^+^ T cells. (B) Polyfunctional CD4^+^ T cells (IFNγ + TNFα^+^ IL2^+^ CD4^+^ T cells) quantified as a percentage of CD4^+^ T cells. (C) IFNγ +, TNFα^+^ and IL2^+^ CD8^+^ T cells, as a percentage of CD8^+^ T cells. (D) Polyfunctional CD8^+^ T cells (IFNγ + TNFα^+^ IL2^+^ CD8^+^ T cells), as a percentage of CD8^+^ T cells. (**E**) Cytokine production by splenocytes after 3 day ex vivo restimulation with full-length Spike protein. The cytokines quantified (in pg/mL) include Th1 cytokines (IFN, TNFα, IL-2), Th2 cytokines (IL-4, IL-5, IL-13), and IL-6. Data presented as mean SEM with n = 4-5 mice per group; * p<0.05, ** p<0.01 by Kruskal-Wallis test with Dunn’s post-test.

To determine the nature of the immune response generated by our glyco-polymer conjugate platform, we quantified the amounts of various cytokines secreted by splenocytes after three days of restimulation with whole Spike protein. Cells collected from the spleens of animals treated with Spike-p(Man-TLR7) and Spike-p(Man-TLR7)+alum produced significantly more Th1 cytokines, specifically IFNγ and IL-2, compared to most other groups (**Fig. 5E**). Interestingly, we observed that splenocytes from mice vaccinated with Spike-p(Man-TLR7)+alum also secreted increased levels of IL-6 upon restimulation, as well as Th2 cytokines, specifically IL-4 and IL-13, compared to other groups (**Fig. 5E**). Nevertheless, the ratio of IFNγ to IL-4, IL-5, IL-13, and IL-6 was significantly higher for Spike-p(Man-TLR7), as well as Spike-p(Man-TLR7)+alum in some cases, compared to benchmark groups (**Fig. S10A**). Increased secretion of IL-17A was not observed upon restimulation of splenocytes from mice vaccinated with Spike-p(Man-TLR7) or Spike-p(Man-TLR7)+alum (**Fig. S10B**). At the same time, all adjuvanted groups showed some elevation in IL-10 secretion (**Fig. S10B**). In summary, we demonstrated that our Spike-p(Man-TLR7) platform induces strong functional Th1-biased T cell responses.

## DISCUSSION

The COVID-19 pandemic has resulted in a global health, economic, and social crisis requiring a rapid response from researchers around the world to develop an effective vaccine against the virus. That urgency led to successful clinical trials and the emergency-use authorization of several vaccine candidates globally, although the considerable uncertainties and failure rates inherent in the process are highlighted by high-profile dropouts, as seen in the case of Sanofi/GSK’s and Merck’s vaccine candidates. Therefore, the global vaccination effort has focused on putting forth as many candidates as possible, in the event that any of the frontrunners failed to live up to their promise in preceding development stages. Based on our recent successful deployment of a synthetic glyco-polymer-based vaccine in addressing the difficulties in the field of malaria vaccination (*42*), we adapted the vaccine platform for SARS-CoV-2 by conjugating SARS-CoV-2 viral proteins to the DC-targeted, TLR7 agonist-containing polymer p(Man-TLR7).

Since the beginning of the COVID-19 pandemic, we have learned that the natural responses to SARS-CoV-2 are temporary and decline quickly after recovery. Generation of long-lasting neutralizing antibody responses to components of SARS-CoV-2 is a primary goal of vaccines that would prevent or limit the infection’s severity. The Spike protein has been one of the major antigens used in vaccines to elicit potent antibody responses, with some formulations focusing on the full-length Spike protein and others focusing on only the RBD domain of Spike. As such, after successfully formulating two bioactive conjugates, RBD-p(Man-TLR7) and Spike-p(Man-TLR7), we investigated the humoral responses generated in vaccinated mice using a prime-boost regimen. Both RBD-p(Man-TLR7) and Spike-p(Man-TLR7) resulted in high levels of IgGs against the corresponding immunogen. However, with our platform, only Spike-p(Man-TLR7) elicited neutralizing antibodies that exceeded the FDA-recommended VNT. Because RBD is a smaller antigen than the full-length Spike protein, conjugation to p(Man-TLR7) could potentially mask epitopes important for eliciting neutralizing antibodies. In fact, multiple lysine residues available for conjugation are located near the ACE2 binding site on RBD (*7, 8*). Because the conjugation occurs randomly on sterically-accessible lysine residues, and Spike is a larger protein with more lysine residues available for conjugation, it is statistically less likely for the p(Man-TLR7) polymer to completely mask this site on Spike. Additionally, since there are neutralizing epitopes on Spike outside of the RBD region (*12*), vaccines using the full-length Spike protein have the potential to elicit neutralizing antibodies against a broader range of epitopes.

Our studies also sought to determine how the combinatorial application of adjuvants may modulate the anatomic localization and efficacy of immune responses. In particular, we explored the use of the depot-forming adjuvant alum, which is commonly seen in other clinical-stage vaccines and COVID-19 vaccine candidates (*45–47, 59*). We observed that alum seemed to have an effect on the anatomic localization of germinal center responses and therefore antibody generation. Vaccination with Spike-p(Man-TLR7) resulted in higher numbers of GC B cells in the spleen, whereas vaccination with Spike-p(Man-TLR7) in combination with alum resulted in higher numbers of GC B cells in the draining lymph nodes (**Fig. 4, A and B**). These findings suggest that alum retains vaccines near the site of injection, whereas in its absence, Spike-p(Man-TLR7) can diffuse more systemically. This is in agreement with one of the proposed mechanisms of action for alum being the formation of an antigen depot that results in the slow drainage of antigen from the injection site (*55–57*). This slow drainage has important implications for vaccine efficacy, as it has been shown previously that controlled prolonged release of antigen can greatly enhance humoral responses upon vaccination (*66*). Although the depot effect of alum has been called into question (*67*), it is nevertheless consistent with our observations of improved magnitude, breadth, and neutralization activity of the humoral responses elicited by Spike-p(Man-TLR7)+alum, and may be worth a further investigation outside the scope of this study.

Both CD4^+^ and CD8^+^ T cells play an important role in the prevention and mitigation of SARS-CoV-2 infection (*65, 68*). Evidence suggests that patients who recovered from COVID-19 had relatively high T cell levels compared to patients who had severe disease complications and died (*69, 70*). Notably, T cell responses are more durable than antibody responses which points to their importance in establishing long term protection against the virus (*32*). Unfortunately, most reported COVID-19 vaccine candidates induce low T cell responses in mice. In contrast, we observed that immunization with Spike-p(Man-TLR7) with and without alum induced a large fraction of antigen-specific polyfunctional CD4^+^ T cells, much higher than that observed for clinical benchmark formulations. Surprisingly, Spike-p(Man-TLR7)+alum also induced large numbers of antigen-specific CD8^+^ T cells. Although alum is often considered a poor CD8^+^ T cell adjuvant (*71, 72*), studies have shown that antigen-specific CD8^+^ T cells can be generated after vaccination with antigen and alum (*73*).

A preferential Th1-biased immune response, as opposed to a Th2-biased response is also desirable from COVID-19 vaccines (*38–40, 74, 75*). Here, a Th1 bias was observed in both the IgG isotypes (an increased IgG2b:IgG1 ratio, **Fig. 2, C and D**) and cytokines secreted from splenocytes harvested from mice vaccinated with Spike-p(Man-TLR7) (**Fig. 5, C and D; Fig. S9B**). While alum synergistically enhances some of the humoral responses seen with Spike-p(Man-TLR7) vaccination, it decreases the Th1-biased responses observed. This is unsurprising, as alum is known to preferentially induce a Th2 response (*76*). As such, when deciding whether to use our p(Man-TLR7) platform alone or in combination with alum, this balance between favorable humoral responses and a skewing away from a Th1 bias must be considered.

There are several translational advantages to our approach for an effective next-generation vaccine. First, the conjugation strategy employed can be performed on any amine-containing antigen, including whole proteins and peptides. This means that, as viral protein mutations emerge, our platform can be easily adapted, and the conjugation can be conducted on newly identified variants. In addition, the APC-directed components of p(Man-TLR7), mannose and the TLR7 agonist, are universally advantageous across species. The pattern recognition receptors recognizing mannose residues are expressed by APCs in both mice and humans and have been shown to play a significant role in antigen capture and processing (*77–79*), while the TLR7 agonist can also be swapped for other TLR agonists. This is particularly pertinent in human immunology, as not only TLR7 (present mostly in plasmacytoid DCs) but also Toll-like receptor 8 (TLR8, expressed by myeloid DC, monocytes and monocyte-derived DCs) agonists have been shown to be necessary to drive strong B and T cell-mediated immune responses (*80*).

Despite the advantages of the flexible antigen-adjuvant conjugation chemistry, this is also the source of the primary limitation of the p(Man-TLR7) vaccine platform. Although the conjugation of antigen to the polymer is via a self-immolative linker, we observed that this may lead to reduced activity of the antigen, in terms of recognition of its target receptor (**Fig 1C and Fig. S3C**). This suggests that the smaller the antigen, the fewer the number of potential epitopes, and the higher the chances that conjugation to p(Man-TLR7) may lead to steric blockade of the receptor-binding site. This may adversely affect the quality of resultant antigen-specific humoral responses, as seen in the case of RBD-p(Man-TLR7) (**Fig. S4**). This is because intracellular processing in the endosomes of APCs is required for release of the antigen from the rest of the construct, whereas humoral responses are partly dependent on extracellular interactions with B cell receptors in the germinal centers of secondary lymphoid organs. For most antigens, however, this will not be a relevant issue, and it may be possible to optimize the polymer to protein ratio if this issue does arise.

In conclusion, to address a global need for next-generation vaccines against SARS-CoV-2, we have developed the Spike-p(Man-TLR7) vaccine platform and demonstrated its efficacy in mice. We found that conjugating the Spike protein to our polymeric glyco-adjuvant improves Spike’s immunogenicity through inducing both potent neutralizing humoral and high-quality cellular responses. We demonstrated that Spike-p(Man-TLR7) is efficacious in elderly mice, and antibody responses are long-lasting. In addition, we determined that combining Spike-p(Man-TLR7) with alum further enhanced immune responses, often exceeding those elicited by mimics of clinical-stage vaccine candidates. Whether in the global fight against SARS-CoV-2 or another pathogen, these studies highlight the adaptability of the modular p(Man-TLR7) platform and reinforce the translational potential of our polymeric glyco-adjuvant to be used in next-generation vaccines.

## MATERIALS AND METHODS

### Study design

This study was designed to test the immunogenicity of an APC-targeting vaccine platform consisting of either prefusion-stabilized Spike protein or its RBD, conjugated to the polymeric glyco-adjuvant p(Man-TLR7). The goal was to develop a next-generation vaccine platform in response to the ongoing COVID-19 pandemic. In the study, the humoral response in mice vaccinated with Spike-p(Man-TLR7), Spike-p(Man-TLR7)+alum, or RBD-p(Man-TLR7) was characterized by evaluating the antibody titers (IgG and IgA) via ELISA, as well as through a viral peptide array and virus neutralization assay. The lymphocyte responses were characterized by flow cytometry, and B and T cellular reactivity were assessed by quantification of antibody or cytokine expression following antigen restimulation. The studied platform’s immunogenicity was compared to that of the following clinically relevant vaccine formulations: Spike+AS04-L, Spike+AS03-L and Spike+alum. Statistical methods were not used to predetermine necessary sample size, but sample sizes were chosen on the basis of estimates from pilot experiments and previously published results such that appropriate statistical tests could yield statistically significant results. All experiments were replicated at least twice except for Figs. 2I, 2J, 3, as well as the experimental groups Spike+AS03-L, Spike+alum, and Spike-p(Man-TLR7)+alum (once). In animal studies, all mice were treated in the same manner. Animals were randomly assigned to a treatment group, and analyses were performed in a blinded fashion. Production of the studied conjugates was performed multiple times to ensure reproducibility. Samples were excluded from analysis only when an animal developed a health problem for a nontreatment-related reason, according to the animal care guidelines. Statistical methods are described in the “Statistical analysis” section.

### Animals

All studies with animals were carried out in accordance with procedures approved by the Institutional Animal Care and Use Committee at the University of Chicago (protocol # 72551) and housed in a specific pathogen-free environment at the University of Chicago. C57Bl/6 female mice aged 8, 21, 47 or greater than 64 weeks were obtained from The Jackson Laboratory.

### Synthesis and characterization of p(Man-TLR7) polymer

The polymeric glyco-adjuvant p(Man–TLR7) was synthesized via a reversible addition– fragmentation chain transfer (RAFT) polymerization using an azide-modified RAFT agent, a biologically inert comonomer (N-(2-hydroxypropyl) methacrylamide, HPMA) and two functional monomers: one synthesized from D-mannose, and the other from a potent TLR7 ligand (mTLR7) (**Fig. S2A**), as described previously (*42*). Molecular weight and polydispersity of the p(Man–TLR7) construct were measured by size exclusion chromatography (molecular weight target at ~20 kDa), and was composed of a 1:2.1:3.5 molar ratio of mTLR7:mannose monomer:HPMA, as measured by mTLR7-specific UV absorbance and ^1^H NMR.

### Sodium dodecyl sulfate-polyacrylamide gel electrophoresis (SDS-PAGE)

SDS-PAGE was performed on stain-free 4-20% gradient gels (Bio-Rad). Samples run under reducing conditions were incubated for 15 min at 95°C with 710 mM 2-Mercaptoethanol. After electrophoresis, gel images were acquired with the ChemiDoc XRS+ system (Bio-Rad).

### Spike and RBD protein production

Plasmids encoding (His)_6_-tagged pre-fusion stabilized Spike protein or RBD protein (sequences in **Table S1**) were obtained from the laboratory of Florian Krammer (Mount Sinai School of Medicine, New York, NY). Suspension-adapted HEK-293F were maintained in serum-free FreeStyle 293 Expression Medium (Gibco). On the day of transfection, cells were inoculated into at a concentration of 1×10^6^ cells/mL. 1 mg/mL plasmid DNA was mixed with 2 mg/mL linear 25 kDa polyethyleneimine (Polysciences) and transfected in OptiPRO SFM medium (4% final volume). After 7 days of culture, supernatants were harvested, and purification was performed as described previously (*81*). Purified proteins were tested for endotoxin via HEK-Blue TLR4 reporter cell line (Invivogen, San Diego, CA) and endotoxin levels were confirmed to be less than 0.01 EU/mL. Protein purity was assessed by SDS-PAGE as described previously (*81*). Protein concentration was determined through absorbance at 280 nm using NanoDrop (Thermo Scientific).

### Surface plasmon resonance (SPR) measurements

SPR measurements were made using a Biacore X100 SPR system (Cytiva, Marlborough, MA). At the beginning of each cycle, 2 μg mL^−1^ recombinant human ACE2-Fc (Sino Biologicals, Beijing, China) in running buffer (0.01 M HEPES pH 7.4, 0.15 M NaCl, 0.005% v/v Surfactant P20) was flowed over a Protein A coated sensor chip (Cytiva) at a flowrate of 5 μL min^−1^ for 780 seconds, resulting in ~700-1100 resonance units corresponding to ligand coating. Spike or RBD protein was then flowed at decreasing concentrations (ranging from 250 nM to 3.9063 nM) in running buffer for contact time of 180 seconds at 30 μL min^−1^, followed by running buffer for a dissociation time of 300 seconds. At the end of each cycle, the sensor chip surface was regenerated with two 30-second pulses of 10 mM glycine pH 1.5 at 30 μL min^−1^. Specific binding of Spike and RBD proteins to ACE2 was calculated by comparison to a non-functionalized channel used as a reference. The experimental results were fitted with Langmuir binding kinetics using the BIAevaluation software (Cytiva, version 2.0.2.).

### Production of RBD-p(Man-TLR7) conjugate

RBD was mixed with 5 molar equivalents of 2 kDa self-immolative PEG linker in a phosphate buffer (pH 7.7) and reacted for 1 hour in an endotoxin-free Eppendorf tube mixing at RT. The reaction solution was then purified via Zeba spin desalting columns with 7 kDa cutoff to remove unreacted linker (Thermo Fisher). Successful linker conjugation was confirmed using gel electrophoresis and comparison to a size standard of the unmodified RBD. RBD-linker construct in PBS (pH 7.4) was then reacted with 30 fold molar excess of p(Man-TLR7) polymer in an endotoxin-free Eppendorf tube for 2 hours, mixing, at RT. Conjugation was confirmed via gel electrophoresis, and conjugates were stored at 4°C.

### Production of Spike-p(Man-TLR7) conjugate

Spike was mixed with 10 molar equivalents of 2 kDa self-immolative PEG linker in a phosphate buffer (pH 7.7) with 0.1% Tween 80 (Sigma) and reacted for 1 hour in an endotoxin-free Eppendorf tube mixing at RT. The reaction solution was then purified via Zeba spin desalting columns with 7 kDa cutoff to remove unreacted linker (Thermo Fisher). Successful linker conjugation was confirmed using gel electrophoresis and comparison to a size standard of the unmodified Spike. Spike-linker construct in PBS (pH 7.4) was then reacted with 30 fold molar excess of p(Man-TLR7) polymer in an endotoxin-free Eppendorf tube for 2 hours, mixing, at RT. Conjugation was confirmed via gel electrophoresis, and conjugates were stored at 4°C.

### Determination of TLR7 content in p(Man-TLR7) conjugates

To determine the concentration of TLR7 content in the polymer and RBD- or Spike- polymer conjugates, the absorbance at 327nm was measured. Known quantities of TLR7 monomer in saline were measured (n=3 independent samples) at 327nm in several concentrations ranging from 8 mg/mL to 1 mg/mL to calculate a standard curve as previously published (*42*). The determined standard curve [TLR7 (mg/mL) = 1.9663* A_327_+0.0517] was then used to calculate TLR7 concentration in the prepared p(Man-TLR7) conjugate.

### Determination of RBD or Spike content in p(Man-TLR7) conjugates

SDS-PAGE was performed as previously stated using a standard curve of RBD or Spike protein and two dilutions of RBD- or Spike-p(Man-TLR7) conjugate samples reduced with 710 mM 2-Mercaptoethanol. Reducing conditions liberate conjugated linker-p(Man-TLR7) from the antigen, allowing for reduced antigen band intensity to be analyzed. The band density of the reduced samples and RBD or Spike standard curve was then analyzed using ImageJ and the RBD or Spike concentration of the samples was calculated using the standard curve generated.

### *In vitro* activity of p(Man-TLR7) conjugates

BMDCs were prepared from C57Bl/6 mice (Jackson Laboratory) as previously described (*82*) and used on day 8–9. For BMDC activation studies, 2×10^5^ cells per well were seeded in round-bottom 96-well plates (Fisher Scientific) in RPMI with 10% FBS and 2% penicillin/streptomycin (Life Technologies), and treated with either Spike or Spike-p(Man-TLR7), then incubated at 37°C. The samples were allowed to culture for 12h at 37°C and cytokine concentration was measured in the media by Ready-Set-Go™ ELISA kits (Thermo Fisher) as detailed in the manufacturer’s instructions.

### ELISA for ACE2 binding

96-well ELISA plates (Nunc MaxiSorp flat-bottom plates, Thermo Fisher) were coated with 10 nM RBD, RBD-p(Man-TLR7), Spike, Spike-p(Man-TLR7), or bovine serum albumin (BSA, Sigma) in PBS overnight at 4°C. The following day, plates were washed in PBS with 0.05% Tween 20 (PBS-T) and then blocked with 2% BSA (Sigma) diluted in PBS for 1 hour at room temperature. Then, wells were washed with PBS-T and further incubated with human ACE2-Fc (Sino Biological) for 2 hours at room temperature. After 6 washes with PBS-T, wells were incubated for 1 hour at room temperature with horseradish peroxide (HRP)-conjugated antibody against human IgG (Jackson ImmunoResearch). After 6 washes with PBS-T, tetramethylbenzidine substrate was added, followed by 10% H_2_SO_4_ after 15 min. Subsequently, the absorbance was measured at 450 nm and 570 nm (Epoch Microplate Spectrophotometer, BioTek).

### Reagents for *in vivo* studies

AS03-like squalene-based adjuvant (AddaS03, InvivoGen), Synthetic Monophosphoryl Lipid A (MPLA, Avanti 699800), and Alhydrogel adjuvant 2% (alum, InvivoGen) were used for vaccination studies. All purchased reagents were used as provided by the manufacturer.

### Vaccination scheme

Mice were vaccinated via s.c. injections into the front two hocks on days 0 and 21. For all vaccine formulations assessed, 10μg of RBD or Spike protein were used. The following amounts of adjuvant were used: 20 μg TLR7 as p(Man-TLR7), 20 μg TLR7 as p(Man-TLR7) + 50 μg alum, 5 μg MPLA + 50 μg alum, 25 μL AS03-L, or 50 μg alum. Excess free p(Man-TLR7) was added to RBD-p(Man-TLR7) and Spike-p(Man-TLR7) conjugates to achieve an exact dose of 20 μg TLR7 per mouse.

### Anti-RBD and anti-Spike antibody analysis

Blood was collected from vaccinated mice weekly or every two weeks into EDTA-K2-coated tubes (Milian). Plasma was separated by centrifugation at 1000 × g for 10 min and stored at − 80°C. Plasma was assessed for anti-RBD or anti-Spike IgGs by ELISA. 96-well ELISA plates (Costar high binding assay plates, Corning) were coated with 10 μg/mL RBD or Spike in 50 mM sodium carbonate/sodium bicarbonate pH 9.6 overnight at 4°C. The following day, plates were washed in PBS with 0.05% Tween 20 (PBS-T) and then blocked with 1× casein (Sigma) diluted in PBS for 1 hour at room temperature. Then, wells were washed with PBS-T and further incubated with various dilutions of plasma for 2 hours at room temperature. After 6 washes with PBS-T, wells were incubated for 1 hour at room temperature with horseradish peroxide (HRP)-conjugated antibody against mouse IgG, IgG1, IgG2b, IgG2c, IgG3, or IgA (Southern Biotech). After 6 washes with PBS-T, bound anti-RBD or anti-Spike antibodies were incubated with tetramethylbenzidine substrate for 18 min. 3% H_2_SO_4_ with 1% HCl was added at that time, and the absorbances at 450 nm and 570 nm were immediately measured (Epoch Microplate Spectrophotometer, BioTek). For all subsequent analysis, the absorbance at 570 nm was subtracted from the absorbance at 450 nm. For titer analysis, the average background plus four times the standard deviation of the background was subtracted from the absorbance values. Titers were calculated as reciprocal dilutions giving values > 0.01. The assay was able to detect titers ranging between 10^−2^ and 10^−7^. An arbitrary value of 0 was assigned to the samples with absorbances below the limit of detection for which it was not possible to detect the titer. For AUC analysis, the fold over the median background absorbance was calculated for each sample, and GraphPad Prism (version 8) was then used to calculate the AUC of the log-transformed plot.

### Antibody epitope breadth determination via peptide array

Antibody specificity to linear epitopes of the Spike protein was analyzed using a CelluSpots™ Covid19_hullB Peptide Array (Intavis Peptide Services, Tubingen, Germany) according to the manufacturer’s protocol. The array comprises 254 peptides spanning the full-length sequence of the Spike protein (NCBI GenBank accession # QHD43416.1), with each 15-mer peptide offset from the previous one by 5 amino acids. Briefly, peptide arrays were blocked in casein blocking solution at 4 °C overnight. Arrays were then incubated with pooled serum diluted 1:200 in blocking buffer for 6 h at room temperature (RT) on an orbital shaker (60 rpm) and then washed 4 times with PBS with 0.05% Tween 20 (PBS-T). Following the fourth wash, arrays were incubated for an additional 2 h at RT and 60 rpm with goat anti-mouse IgG conjugated to HRP (Southern Biotech) diluted 1:5000 in blocking solution. Arrays were washed another 4 times with PBS-T. Spots were detected with Clarity™ Western ECL Substrate (Bio-Rad), and chemiluminescence was measured using a ChemiDoc XRS+ system Gel Documentation System (Bio-Rad). Spots were analyzed using Spotfinder software (version v3.2.1).

### SARS-CoV-2 virus neutralization assay

Heat-inactivated plasma from vaccinated or control mice were serially diluted in DMEM with 2% FBS, 1% Penicillin-Streptomycin, and 10mM Non-Essential Amino Acids (Gibco; mixture of glycine, L-alanine, L-asparagine, L-aspartic acid, L-glutamic acid, L-proline, & L-serine)), and subsequently incubated with 400 plaque-forming units of SARS-CoV-2 virus (strain nCov/Washington/1/2020, provided by the National Biocontainment Laboratory, Galveston TX, USA) for 1 h at 37°C. These mixtures were then applied to Vero-E6 cells and maintained until > 90% cell death occurred in the “no serum” control condition (about 4-5 days). After that, cells were washed with PBS and fixed with 10% formalin, before being stained with crystal violet. Viability was then quantified using a Tecan infinite m200 microplate reader (absorbance 595 nm). Viral neutralization titer represents the greatest plasma dilution at which 50% of SARS-CoV-2-induced cell death is inhibited (EC50). To determine the EC50, data were fit using a least squares variable slope four-parameter model. To ensure realistic EC50 values, we considered a dilution (1/X) of X = 10-1 to be 100% neutralizing and a dilution of X = 108 to be 0% neutralizing and constrained EC50 > 0. Plasma from convalescent human COVID-19 patients were provided by Ali Ellebedy (Washington University School of Medicine, St. Louis, MO; Catalog # NR-53661, NR-53662, NR-53663, NR-53664, and NR-53665).

### Preparation of single cell suspensions from organs

Spleens and injection dLNs were collected on day 28 (7 days post-boost) and stored in ice-cold IMDM (Gibco) until further steps. Spleens were processed into a single-cell suspension via mechanical disruption and passage through a 70μm filter. The splenocytes were washed with PBS and then exposed to ACK lysis buffer (0.155 M NH4Cl, Gibco) for 5 minutes at room temperature to lyse red blood cells. The lymph nodes were mechanically disrupted, then digested at 37°C for 45 min in IMDM with 3.5 mg/mL collagenase D (Roche) before being passed through a 70μm filter. Single cell suspensions were then washed with PBS and resuspended in complete IMDM (IMDM with 10% FBS and 1% Penicillin-Streptomycin).

### Anti-Spike IgG enzyme-linked immunosorbent spot (ELISpot) assay

ELISpot plates (Millipore IP Filter plate) were coated with 20μg/mL Spike in sterile PBS overnight at 4°C. Plates were then blocked using ELISpot Media (RPMI 1640, 1% Glutamine, 10% Fetal Bovine Serum, 1% Penicillin-Streptomycin) for 2 hours at 37°C. Splenocytes from vaccinated mice were seeded in triplicate at a starting concentration of 6.75×10^5^ cell/well and diluted in 3-fold serial dilutions for a total of four dilutions. Plates were incubated for 18 hours at 37°C, 5% CO2 after which the cells were washed off 5× in PBS. Wells were incubated with 100μL IgG-biotin HU adsorbed (Southern Biotech) for 2hr at RT. Next, plates were washed 4× in PBS followed by 100μL HRP-conjugated streptavidin for 1hr at RT. Plates were washed again and incubated with 100μL TMB/well for 5 minutes until distinct spots emerge. Finally, plates are then washed 3× with distilled water and left to dry completely in a laminar flow hood. A CTL ImmunoSpot Analyzer was used to image plates, count spots and perform quality control.

### *Ex vivo* restimulations

Splenocytes were either restimulated *in vitro* with whole Spike protein or Spike peptide pools (PepMix SARS-CoV-2 Spike Glycoprotein, JPT). For Spike protein restimulations, 5×10^5^ cells were incubated with 100 mg/mL Spike protein for 3 days in complete IMDM. After 3 days, the cells were spun down, and the supernatant was used to measure secreted cytokines using a LEGENDplex™ Mouse Th Cytokine Panel kit (BioLegend) according to the manufacturer’s instruction. Approximately 500 events per cytokine was acquired using Attune NxT flow cytometer (ThermoFisher), and analyzed with LEGENDplex v8.0 software.

For Spike peptide restimulations, 2×10^6^ cells were incubated with combined Spike peptide pools (diluted according to manufacturer’s instructions) or equivalent amounts of DMSO (as an unstimulated control) for 6 hours in complete IMDM. After 2 hours of *in vitro* restimulation, GolgiPlug (BD) was added according to manufacturer’s instructions. Cells were then allowed to incubate for 4 more hours before staining for intracellular cytokines and analyzed via flow cytometry, as described.

### Production of RBD protein tetramers

RBD protein expressed with AviTag was purchased from GenScript. Site-specific biotinylation of the AviTag was performed using BirA Biotin-Protein Ligase Reaction kit (Avidity). Next, unconjugated biotin was removed using Zeba spin desalting columns, 7K MWCO (ThermoFisher). The quantification of reacted biotin was performed using the Pierce Biotin Quantification Kit (ThermoFisher). Biotinylated RBD was incubated with either streptavidin-conjugated PE (Biolegend) or streptavidin-conjugated APC fluorophores (Biolegend) for 20 min on ice at a molar ratio of 4:1 of biotin to streptavidin. FITC-labelled Streptavidin (Biolegend) was reacted with excess free biotin to form a non-RBD-specific streptavidin probe as a control. Tetramer formation was confirmed using SDS-PAGE gel. Cells were stained for flow cytometry with all three streptavidin probes at the same time as other fluorescent surface markers at a volumetric ratio of 1:100 for RBD-streptavidin-PE and 1:200 for RBD-streptavidin-APC and biotin-streptavidin-FITC.

### Flow cytometric analysis

The following procedures were all performed at 4°C in the dark. Prepared cells were stained for viability using fixable dyes (Fixable Viability Dye eFluor455, Invitrogen 65-0868-14; Live/Dead Violet Dead Cell Stain Kit, Invitrogen L34964; Fixable Viability Dye eFluor780, Invitrogen 65-0865-14) at 1:500 dilution in PBS with anti-CD16/32 included (1:100 dilution) for 15 minutes. Surface staining was performed in Brilliant Stain buffer (BD Biosciences) using the made in-house tetramers and monoclonal antibodies against the murine targets (**Table S2**). All antibodies and tetramers were titrated to determine optimal working dilutions which often was 1:100 or 1:200. Cells were incubated with the surface stain cocktail for 20 minutes before washing in PBS and fixation. Fixation was performed using the following buffers: for assays without intracellular staining, cells were fixed for 20 minutes using a 2% paraformaldehyde solution; for assays with transcription factor staining, cells were fixed and permeabilized using the Invitrogen FoxP3/Transcription factor kit (eBioscience) according to manufacturer instructions; for assays which required non-transcription factor internal staining (cytokines alone) fixation and permeabilization was performed using the Cytofix/Cytoperm kit (BD Biosciences) according to manufacturer instructions. Assays requiring intracellular staining were performed using antibodies against the murine targets at 1:200 dilution in the corresponding kit permeabilization buffer, according to manufacturer instructions (**Table S2**).Following fixation and/or intracellular staining, cells were resuspended in FACS buffer (PBS pH 7.4 with 2mM EDTA and 2% FBS, made in house) prior to flow cytometric analysis.

### Statistical analysis

Statistical analysis was performed using GraphPad Prism v8. Multiple group comparisons used one-way ANOVA with Tukey’s post-hoc correction, Brown-Forsythe ANOVA with Dunnet’s T3 post-test, or two-way ANOVA with Tukey’s multiple comparisons test. For nonparametric data, the Kruskal-Wallis test, followed by a Dunn’s multiple comparison test, was used. For single comparisons to a specific value, a one-sample *t*-test or Wilcoxon signed rank test was used. Data are presented as mean ± SEM, unless otherwise noted. The n values used to calculate statistics are indicated in figure legends. Significance is indicated as follows, unless otherwise noted: * p<0.05, ** p<0.01, and *** p<0.001.

## Acknowledgements

We acknowledge helpful discussions with Patrick C. Wilson, Jenna J. Guthmiller, Anne I. Sperling, and Aaron Esser-Kahn (University of Chicago, Chicago, IL) and with Robert Baker and David Boltz (Illinois Institute of Technology Research Institute, Chicago, IL) that were instrumental to experimental planning and model development. We acknowledge Suzana Gomes and Tera Lavoie for technical assistance. Parts of this work were carried out at the Cytometry and Antibody Technology Core Facility (Cancer Center Support Grant P30CA014599), the Soft Matter Characterization Facility, the Mass Spectrometry Facility (NSF instrumentation grant CHE-1048528), the Nuclear Magnetic Resonance Facility, the Advanced Electron Microscopy Facility (RRID:SCR_019198), and the Human Immunologic Monitoring Facility (RRID:SCR_017916) at the University of Chicago. We would also like to thank the University of Chicago Animal Resources Center for help and guidance on animal work. We are grateful to the laboratory of Florian Krammer (Icahn School of Medicine at Mount Sinai, New York City, NY) for providing plasmids coding for the Spike RBD, produced with support from the NIH NIAID (Contract # HHSN272201400008C). We are also grateful to the groups of Jesse Bloom (Fred Hutchinson Cancer Research Center, Seattle, WA) and Ali Ellebedy (Washington University School of Medicine, St. Louis, MO) for contributing reagents via the NIH NIAID BEI Resources repository.

## Author contributions

JAH, MAS, and MK oversaw all research; LTG, MMR, TMM, and RW performed the synthesis and characterization of the conjugates; PSB, AM, and EB performed protein expression and purification; BM provided materials; JWR conducted the SPR analysis; all authors designed animal studies. LTG, PSB, TMM, ATA, RPW, LRV, MSS, SC, MN, EAW, AS, NM, RW, and ACT performed animal studies; PSB, VN, KF, SD, and GR designed and performed neutralization assay; LTG, MMR, PSB, TMM, SSY, and ATA wrote the manuscript; all authors proofread the manuscript.

## Data and materials availability

All data are available in the main text or the supplementary materials.

## Supplementary Materials for

**Fig. S1.**
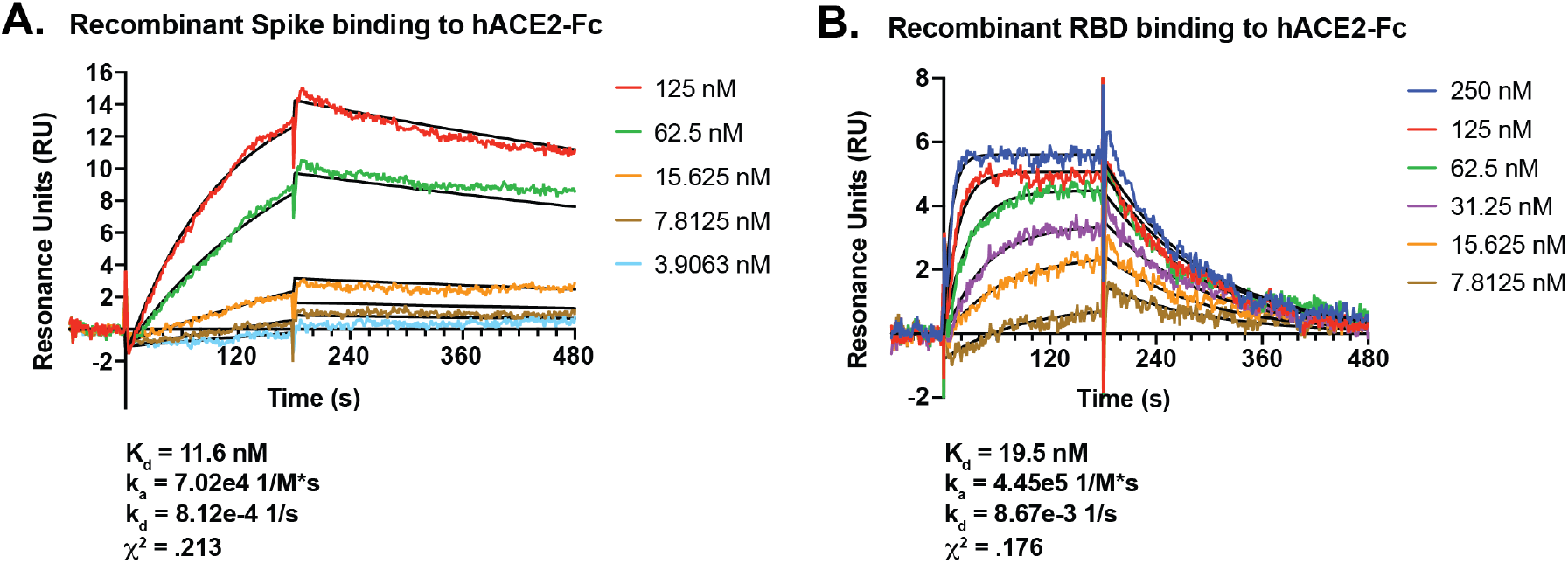
Surface Plasmon Resonance (SPR) analysis of binding of SARS-CoV-2 antigens to human ACE2 (hACE2). Characterization of the binding between (**A**) Spike or (**B**) RBD and hACE2-Fc was conducted using SPR. The graphs represent the real-time binding profile for each antigen and calculated K_d_, k_a_, k_d_ and χ^2^.

**Fig. S2.**
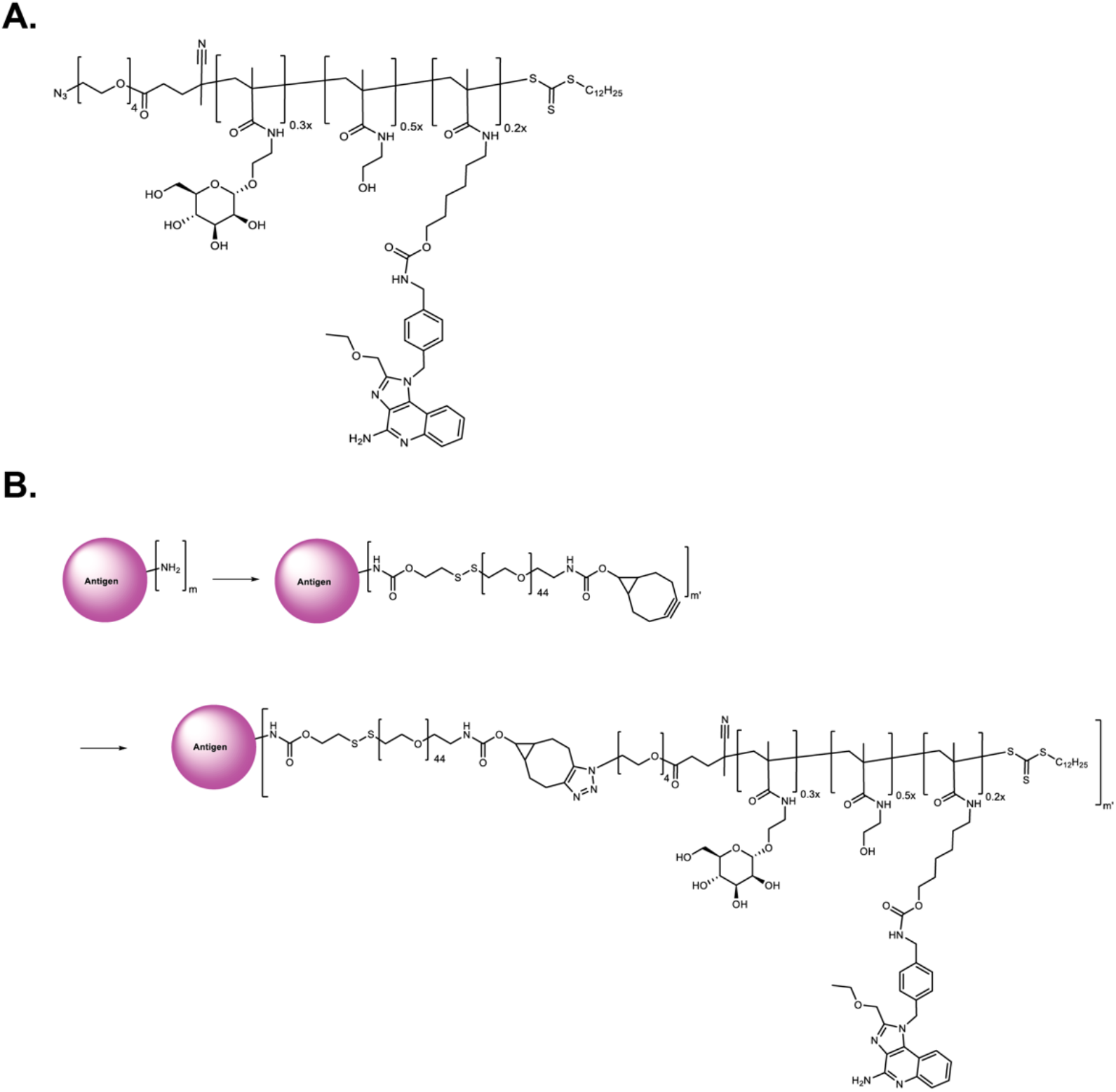
Components of antigen-p(Man-TLR7) platform. (**A**) The polymeric glycol-adjuvant p(Man-TLR7) is a random copolymer composed of monomers with pendant mannose and TLR7 agonist moieties. (**B**) The p(Man-TLR7) polymer is conjugated to amine-containing protein antigens via a two-step conjugation reaction. First, heterobifunctional PEG2000-based linker is attached to the antigen, forming an antigen-linker conjugate. Then, p(Man-TLR7) is reacted with the antigen-linker conjugate via copper-free click-chemistry to form the final antigen-p(Man-TLR7) conjugate.

**Fig. S3.**
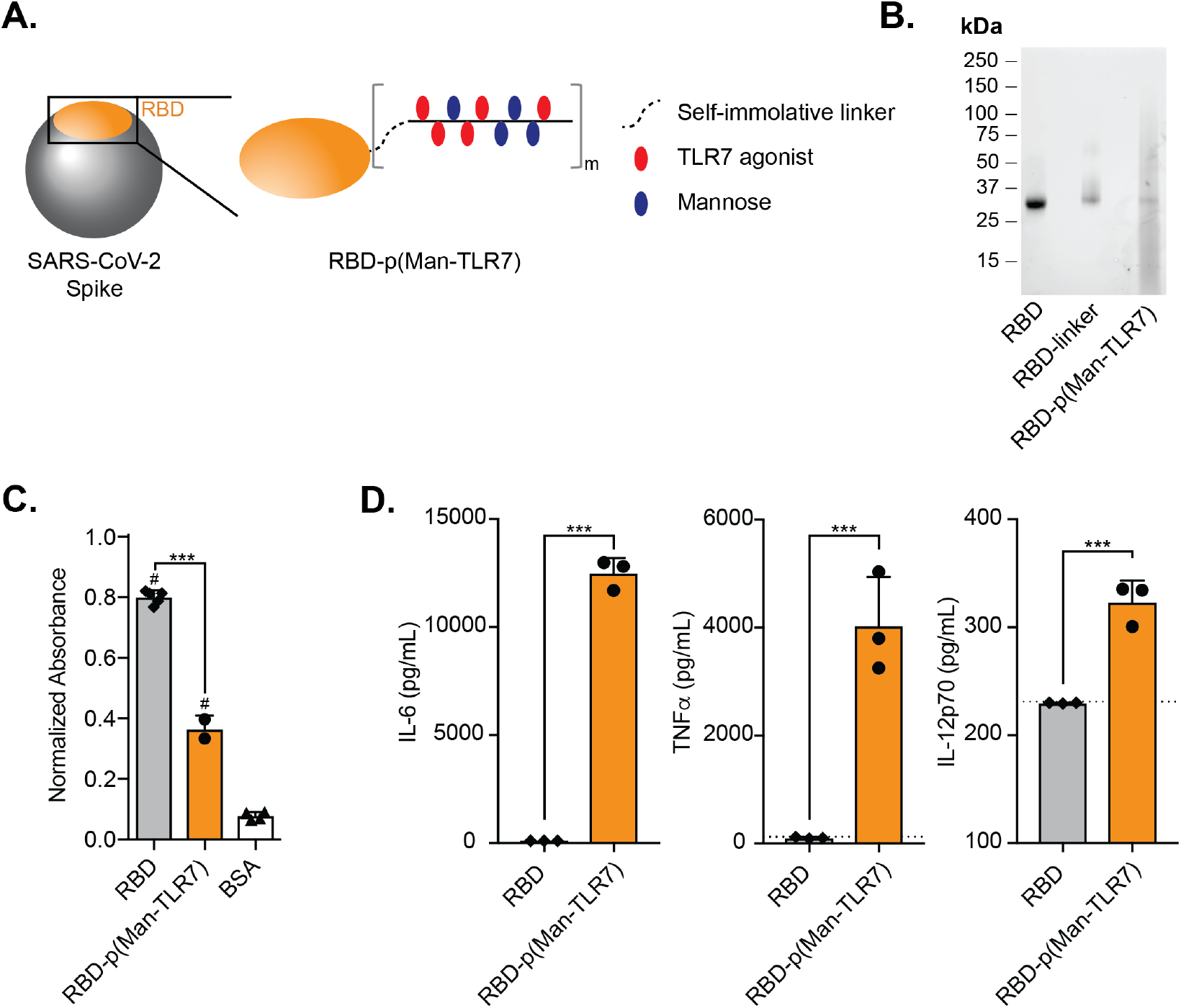
The RBD antigen conjugated to p(Man-TLR7) is a potent activator of BMDCs. (**A**) RBD-p(Man-TLR7) is composed of RBD antigen conjugated, via a self-immolative linker, to a random copolymer synthesized from monomers that either activate TLR7 (red ovals) or target mannose-binding C-type lectins (blue ovals). (**B**) SDS-PAGE analysis of RBD before and after the two step conjugation reaction. The observed band between 15 and 25kDa comes from the free p(Man-TLR7) polymer. (**C**) Analysis of the binding ability of RBD-p(Man-TLR7) to hACE2 via ELISA. (**D**) Concentration of IL-6, TNFα and IL-12p70 in the supernatant of BMDCs stimulated for 18h with either RBD or RBD-p(Man-TLR7) at the concentration corresponding to 25μM of the adjuvant, as determined by ELISA. Dotted horizontal lines represent the assay background. In (C and D), columns and error bars indicate mean+SD; statistical comparisons are based on one-way ANOVA with Tukey’s post-test: *** p<0.001; # p<0.001 as compared to BSA.

**Fig. S4.**
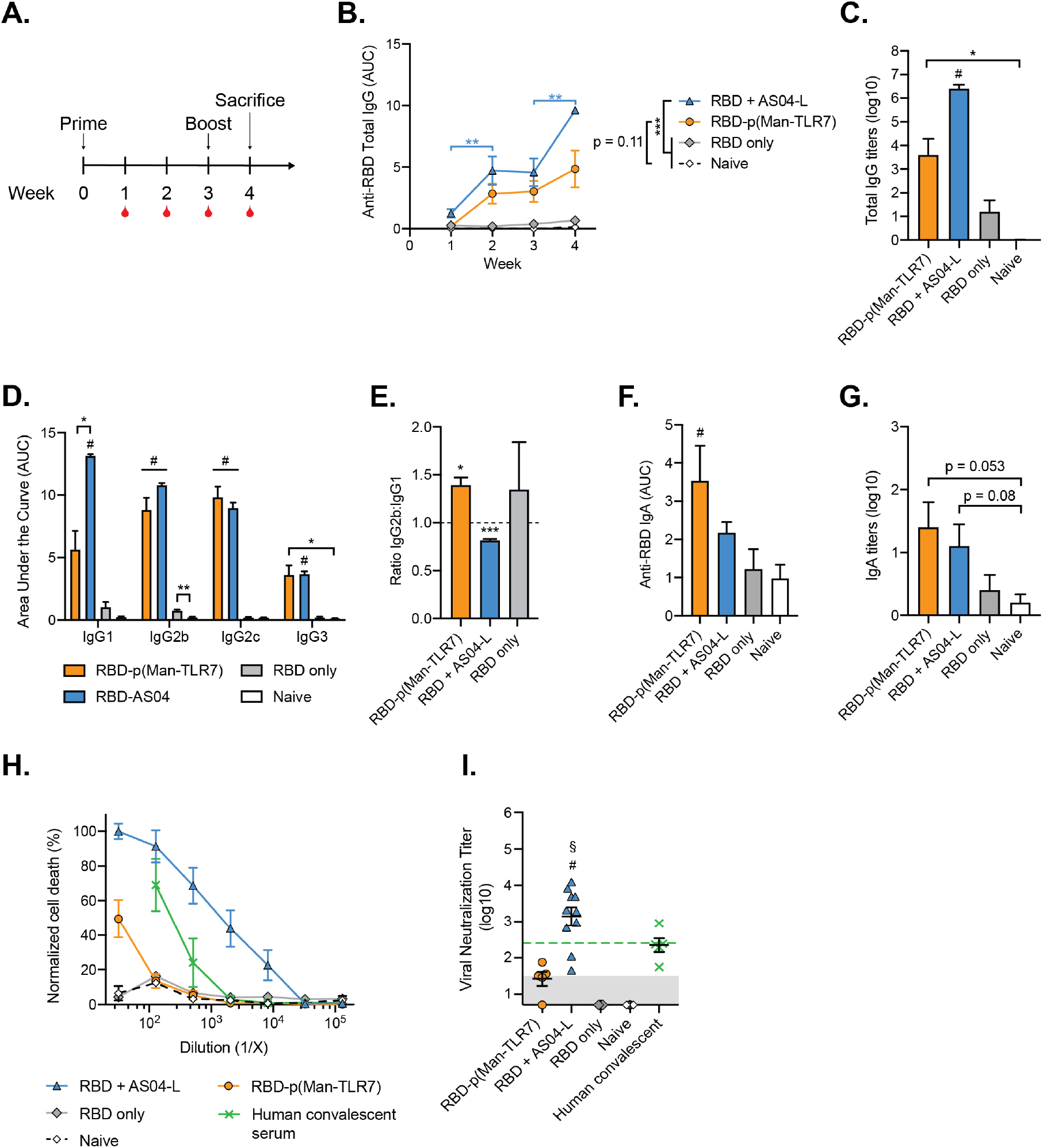
RBD-p(Man-TLR7) vaccination generates RBD-specific antibodies, which fail to potently neutralize SARS-CoV-2. (**A**) Mice were vaccinated with RBD-p(Man-TLR7), RBD+AS04-L, or RBD at weeks 0 (prime) and 3 (boost) and their plasma was collected weekly up until week 4. Plasma from naïve mice was collected at the same time points. (**B**) Total RBD-specific IgG antibodies over time reported as the area under the log-transformed curve (AUC) of absorbance vs. dilution. (**C**) Titers of total RBD-specific IgG antibodies at week 4. (**D**) Comparison of RBD-specific IgG Isotypes (IgG1, IgG2b, IgG2c and IgG3) and (**E**) corresponding IgG2b:IgG1 ratios at one week post boost (week 4). (**F and G**) Circulating anti-RBD IgA antibodies quantified at week 4 via (F) AUC analysis and (G) titration. (**H**) Neutralization assay of live SARS-CoV-2 infection on Vero-E6 cells *in vitro*. SARS-CoV-2 was pre-incubated with plasma isolated from mice at week 4. Percent neutralization was calculated based on viability of cells that did not receive virus (100%) or virus without plasma preincubation (0%). (**I**) Viral neutralization titers, representing plasma dilution at which 50% of SARS-CoV-2-mediated cells death is neutralized. Shaded area represents the lower limit of detection (titer of 2.11); green dotted horizontal line represents FDA recommendation for “high titer” classification (= 2.40). All data are presented as mean±SEM with n = 5-10 mice per group. Comparisons were made using (B) two-way ANOVA with Tukey’s multiple comparison test, (C and D) Brown-Forsythe ANOVA with Dunnet’s T3 test, (E) one sample t test (compared to the theoretical value of 1, representing an unbiased Th1/Th2 response), (F and G) one-way ANOVA with Tukey’s post-test, or (I) Kruskal-Wallis non-parametric test with Dunn’s post-test: * p<0.05, **p<0.01, *** p<0.001; # p<0.05 (for comparison to both RBD and Naïve groups). Additional comparisons were made in (I) using Wilcoxon signed rank test: § p<0.05 (as compared to the FDA “high titer” classification). In (B), comparisons noted on the graph in blue are between the indicated timepoints for RBD+AS04-L, and comparisons noted in the legend are between the indicated groups at week 4. In (B), only relevant statistical comparisons are shown.

**Fig. S5.**
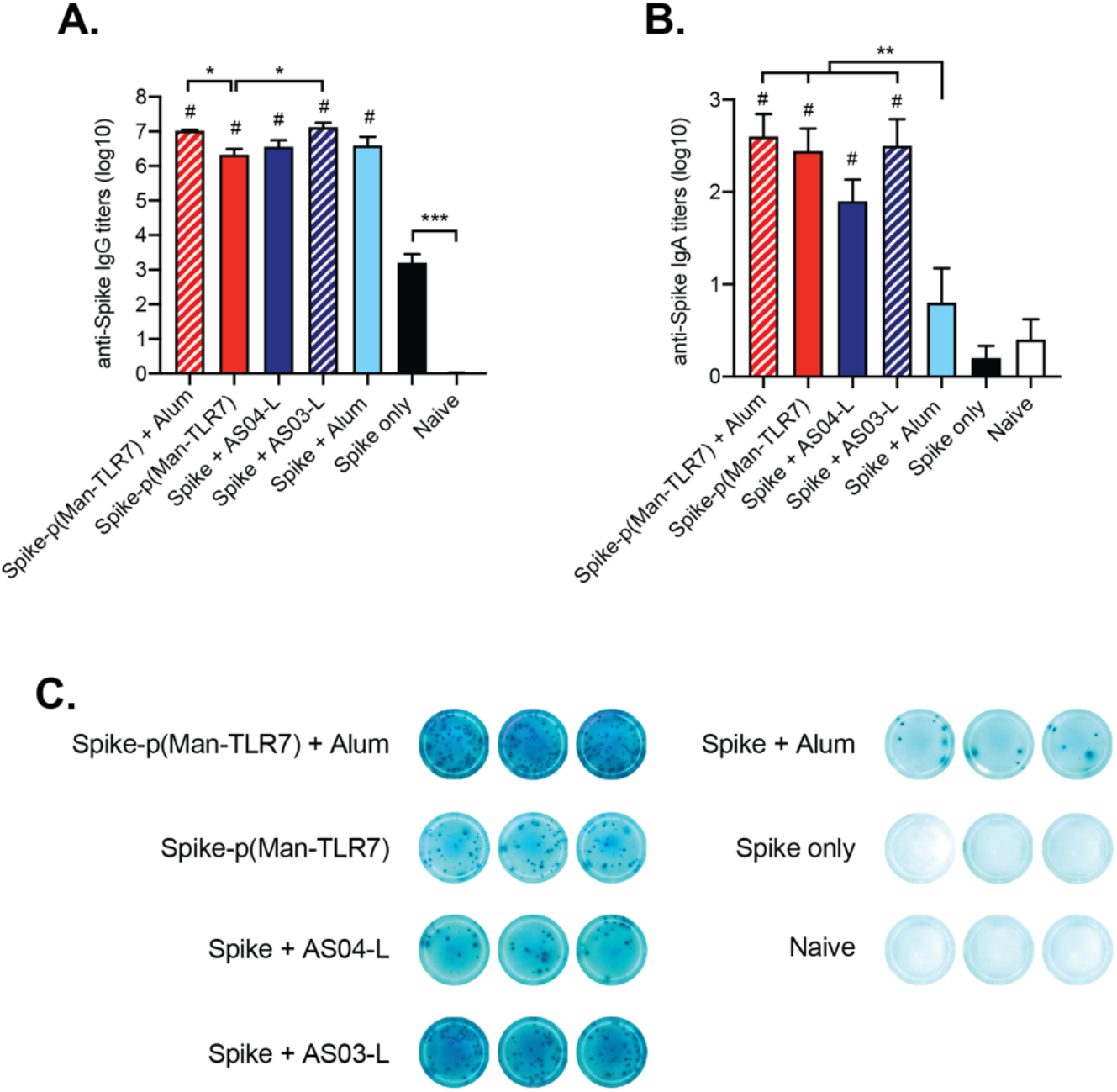
Additional data on Spike-specific humoral responses. (**A**) Total anti-Spike IgG titers and (**B**) IgA titers found one week post-boost in plasma of vaccinated mice (vaccinated as in Fig. 2A). Columns and error bars indicate mean+SEM for n = 4-5 mice per group. Comparisons were made using (A) Brown-Forsythe ANOVA with Dunnett’s T3 post-test or (B) one-way ANOVA with Tukey’s post-test: * p<0.05, **p<0.01 and *** p<0.001; # p<0.05 (for comparison to both Spike and naïve groups). (**C**) Representative ELISpot wells used to quantify Spike-specific IgG^+^ antibody secreting cells from Fig. 1E.

**Fig. S6.**
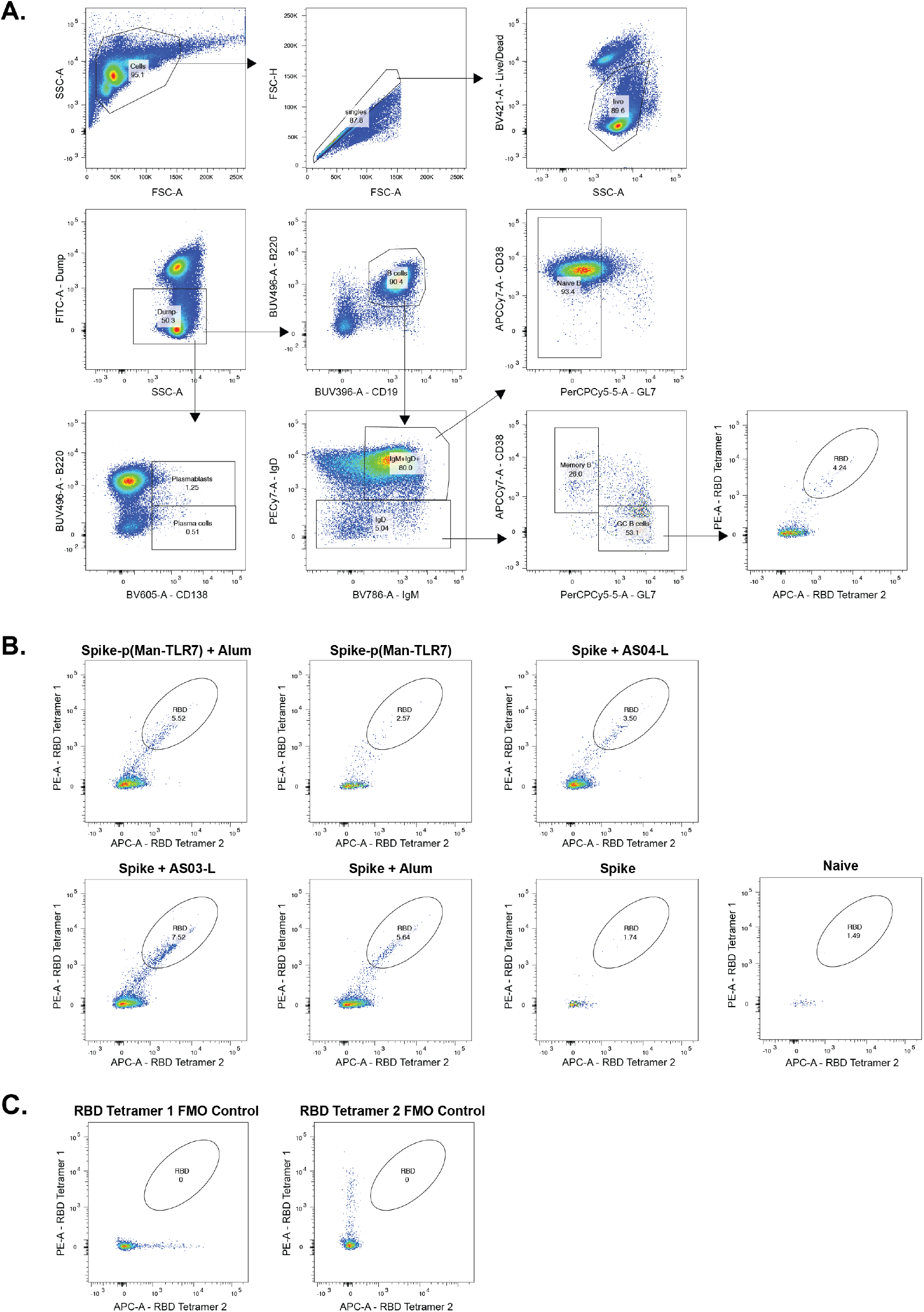
Representative gating used to characterize B cells in Figure 4, A and B. (**A**) Gating strategy used in the analysis of B cells from Fig. 4, A and B. (**B**) Representative flow cytometry plots showing RBD-tetramer^+^ GC B cells from each experimental group. (**C**) Fold minus one (FMO) control flow cytometry plots used for analyzing RBD-tetramer^+^ GC B cells.

**Fig. S7.**
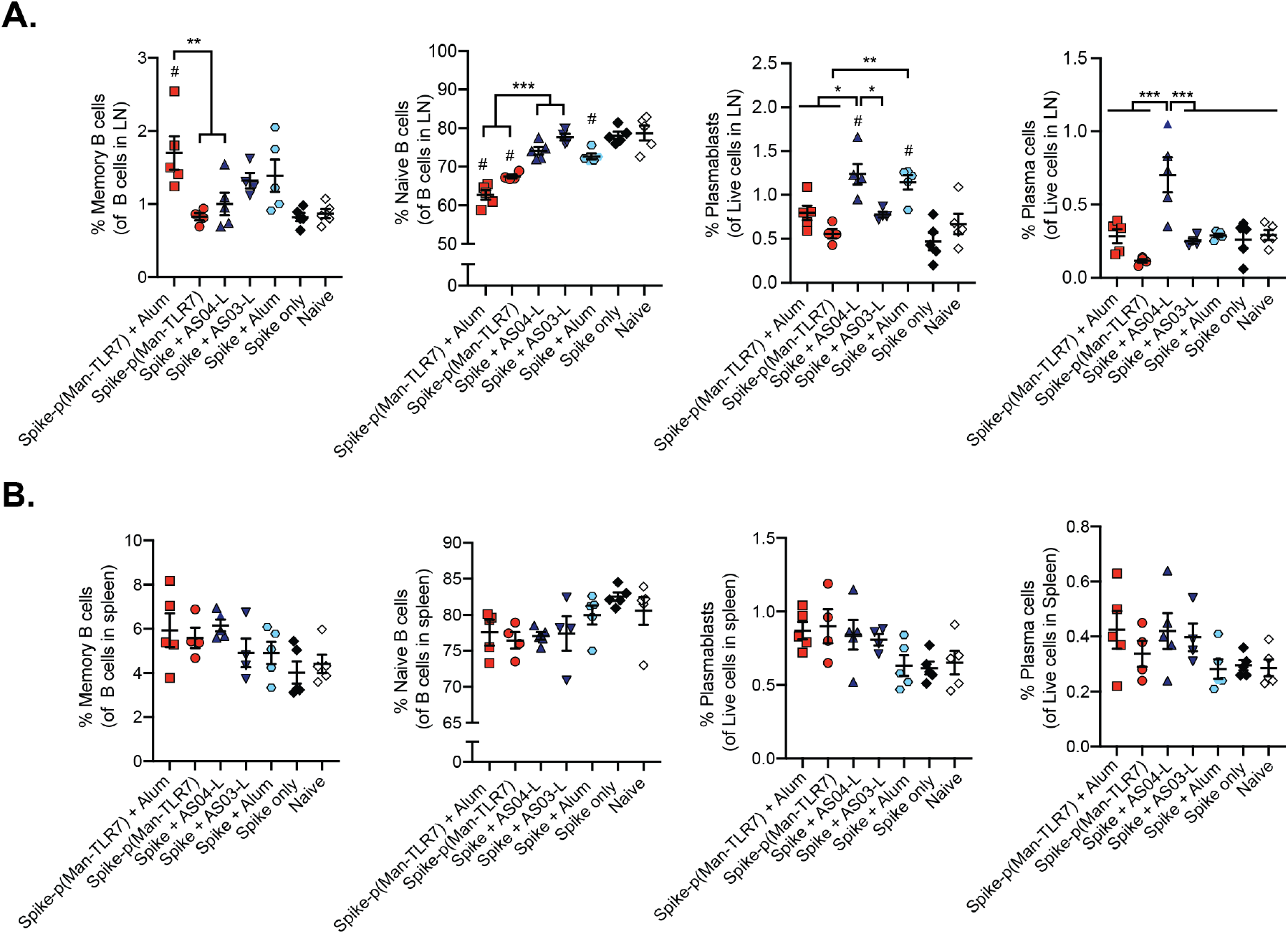
Naïve B cells, memory B cells, plasmablasts, and plasma cells in vaccinated mice 1 week post-boost. (**A and B**) Characterization of B cells resident within the (A) dLNs of the vaccination site or (B) spleen via flow cytometry at week 4 after vaccination (mice were vaccinated as in Fig. 2A). (A-B) From left to right: memory B cells (IgD^+^GL7^−^CD38^+^) within all B cells, naïve B cells (IgM^+^IgD^+^GL7^−^) within B cells, plasmablasts (B220^+^ CD138^+^) within all live cells, and plasma cells (B220^−^CD138^+^) within all live cells. Data presented as mean±SEM with n = 4-5 mice per group; *p<0.05, ** p<0.01, ***p<0.001 via one-way ANOVA with Tukey’s post-test; # p<0.05 as compared to both Spike and naïve.

**Fig. S8.**
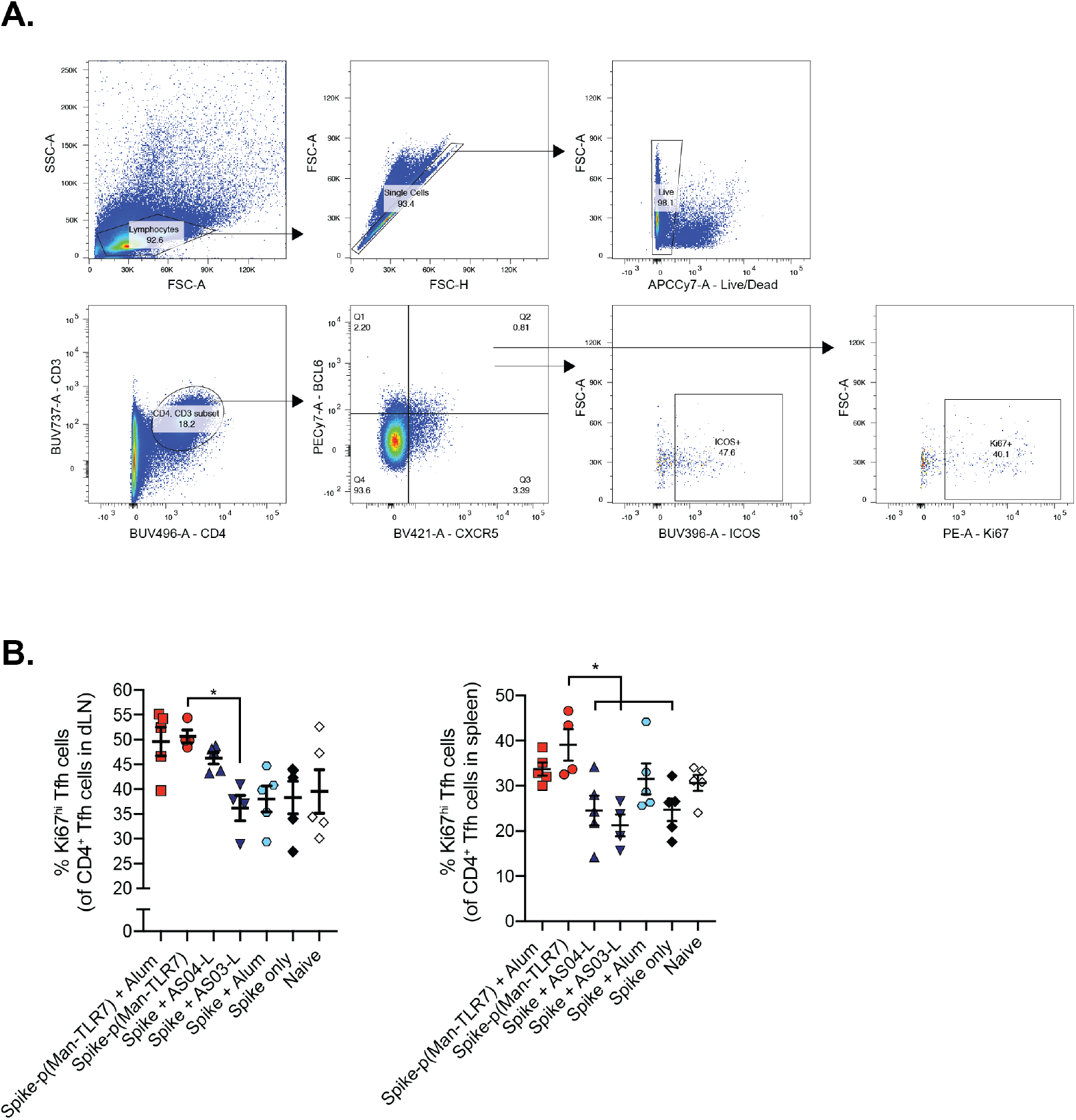
Further characterization of T_fh_ cells in vaccinated mice. (**A**) Gating strategy used for the analysis of T_fh_ cells in Fig. 4C. (**B**) Characterization of proliferating T_fh_ cells (Ki67^hi^ T_fh_ cells, as percentage of all T_fh_ cells) in the dLNs (left) and spleen (right) of vaccinated mice (vaccinated as in Fig. 2A). Data presented as mean±SEM with n = 4-5 mice per group; *p<0.05 by one-way ANOVA with Tukey’s post-test.

**Fig. S9.**
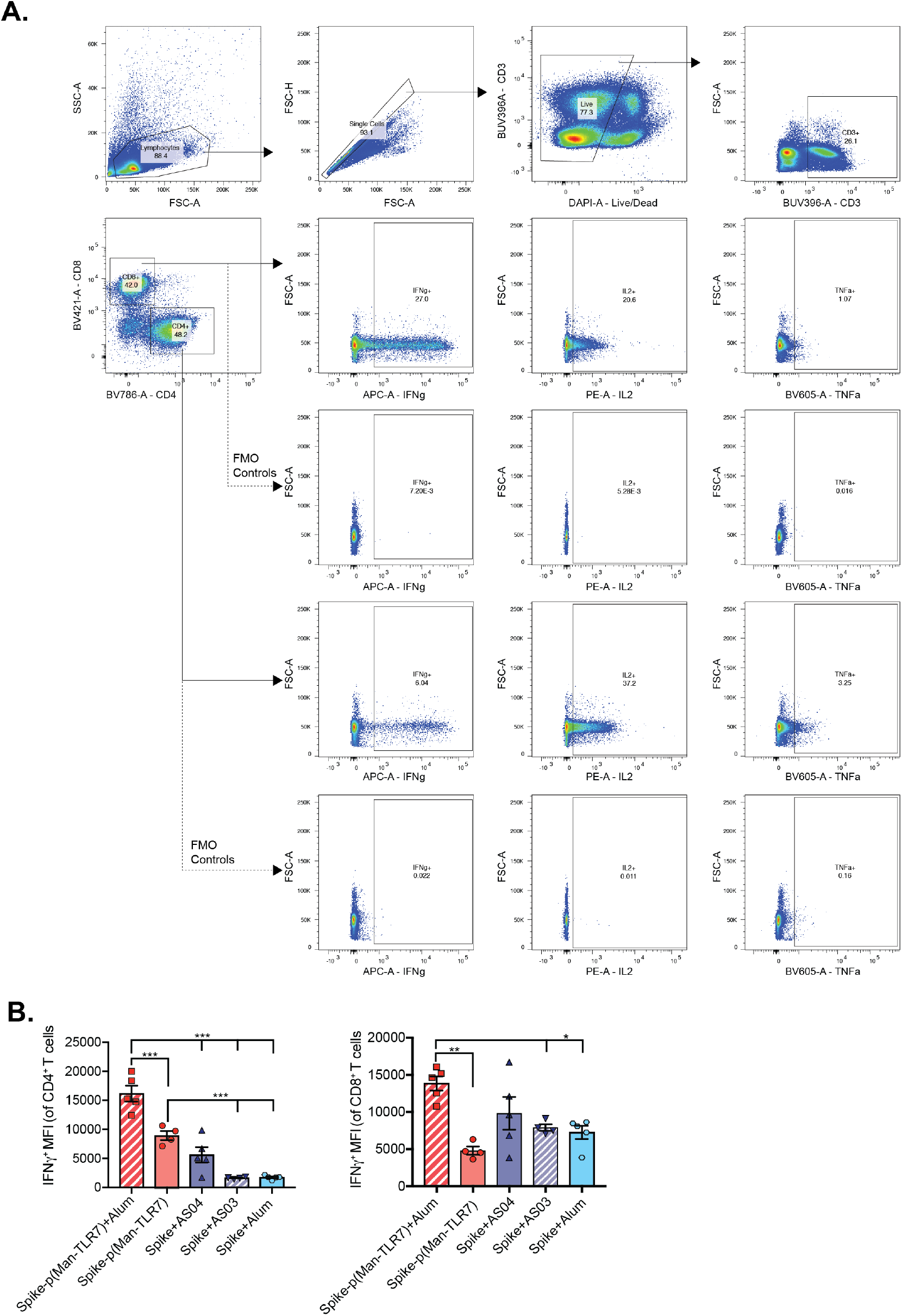
Further characterization of antigen-specific T cells isolated from ex vivo restimulated splenocytes from vaccinated mice. (**A**) Gating strategy used for the analysis of cytokine^+^ T cells from Fig. 5, A to D. (**B**) Mean fluorescence intensity (MFI) of IFNγ + CD4^+^ T cells (left) and IFNγ + CD8^+^ T cells (right). Columns and error bars indicate mean±SEM for n = 4-5 mice per group; *p<0.05, ** p<0.01, ***p<0.001 by one-way ANOVA with Tukey’s post-test.

**Fig. S10.**
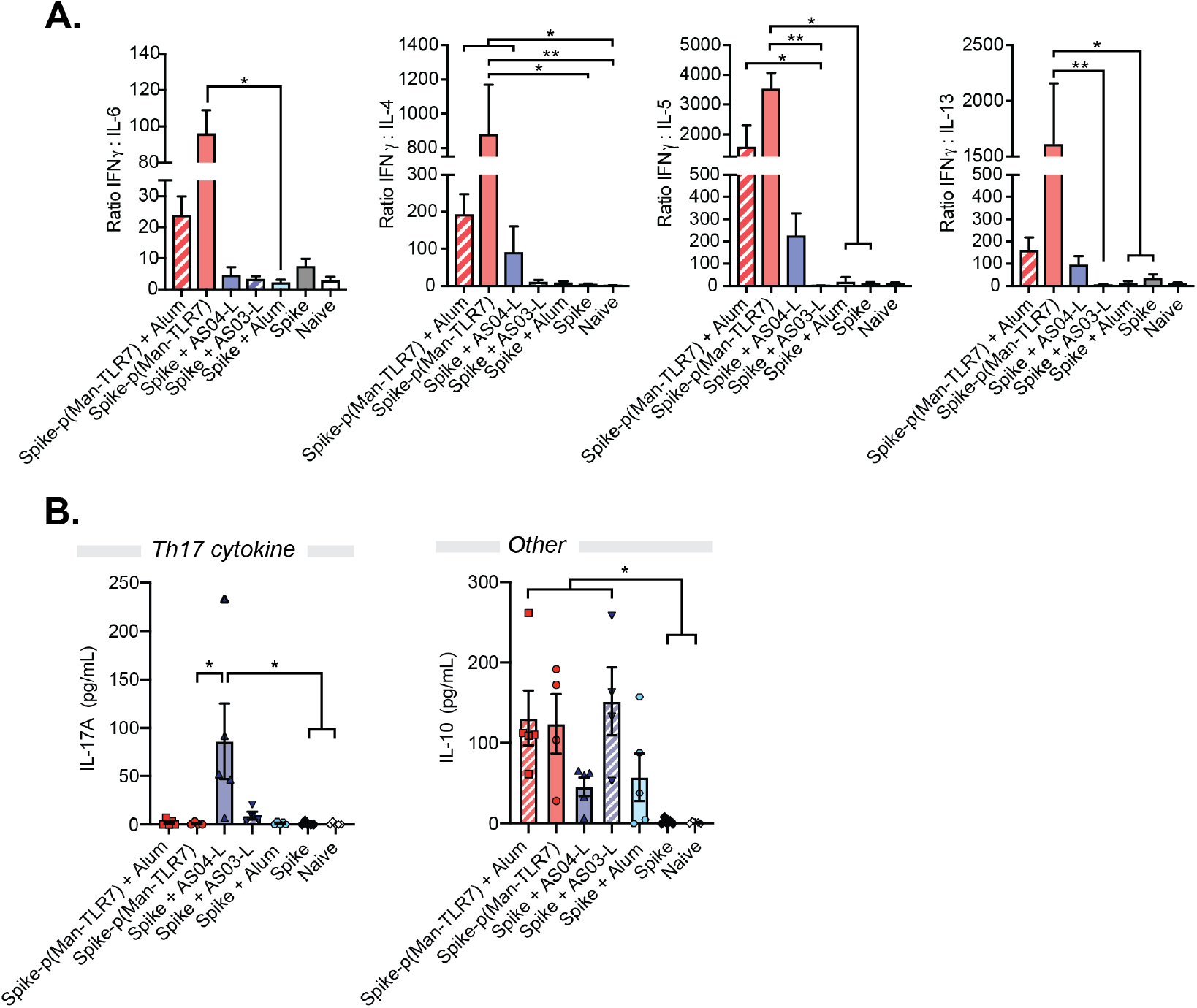
Further characterization of cytokine production upon *ex vivo* splenocyte restimulation with full-length Spike protein. (**A**) Concentration of IL-17A (a T helper cell type 17 (Th17) cytokine) and IL-10 in the supernatant, as quantified by multiplex analysis. (**B**) Ratio (unitless) of IFNγ concentration (pg/mL) to IL-6 and Th2 cytokine (IL-4, IL-5, IL-13) concentration (pg/mL) from Fig. 5E. From left to right, graphs represent the following ratios: IFNγ /IL-6, IFNγ /IL-4, IFNγ /IL-5 and IFNγ /IL-13. Columns and error bars indicate mean±SEM for n = 4-5 mice per group; *p<0.05, ** p<0.01 via Kruskal-Wallis test with Dunn’s post-test.

**Table S1.**
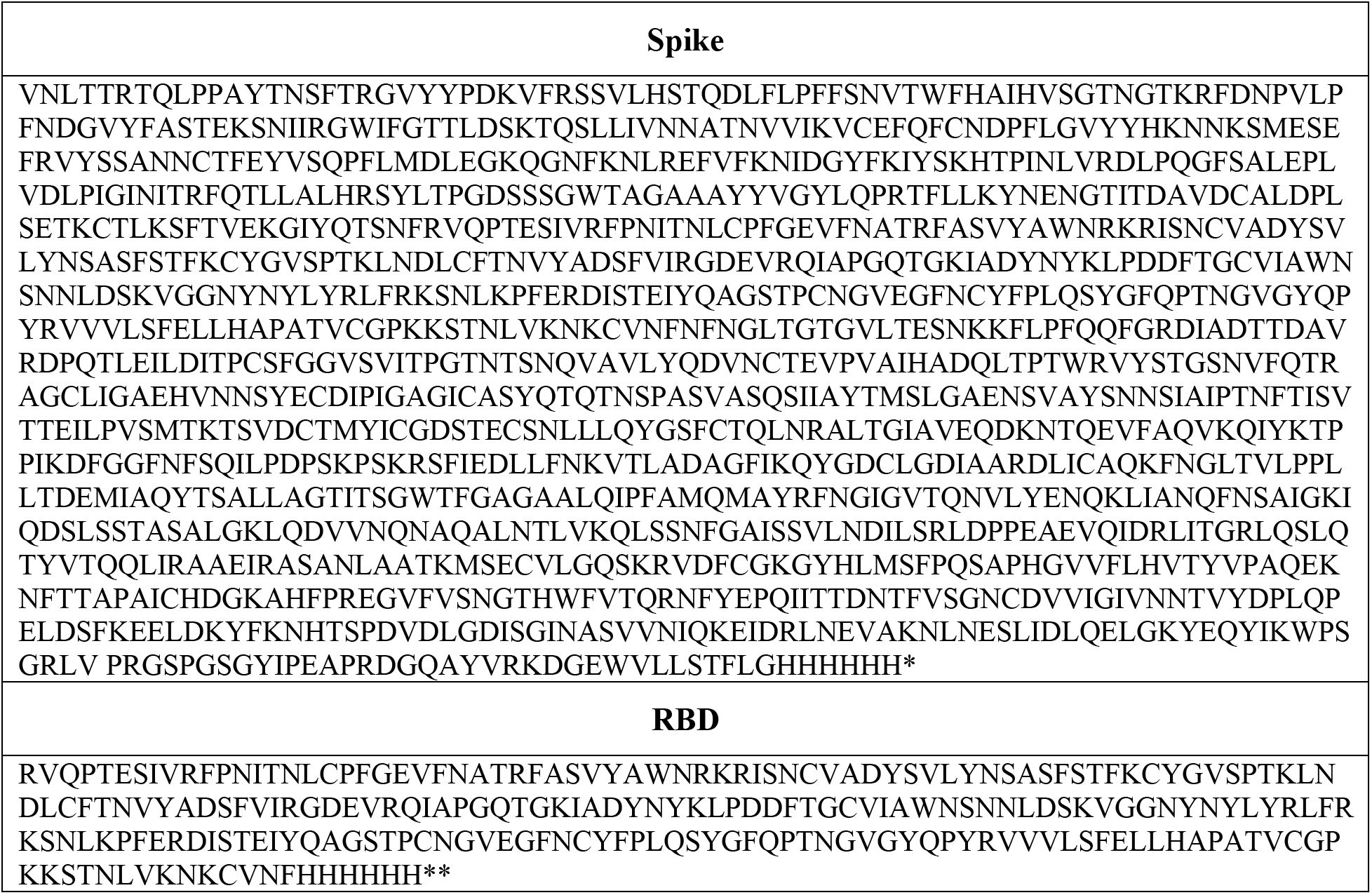
Amino acid sequences of Spike and RBD antigens used in this study.

**Table S2.**
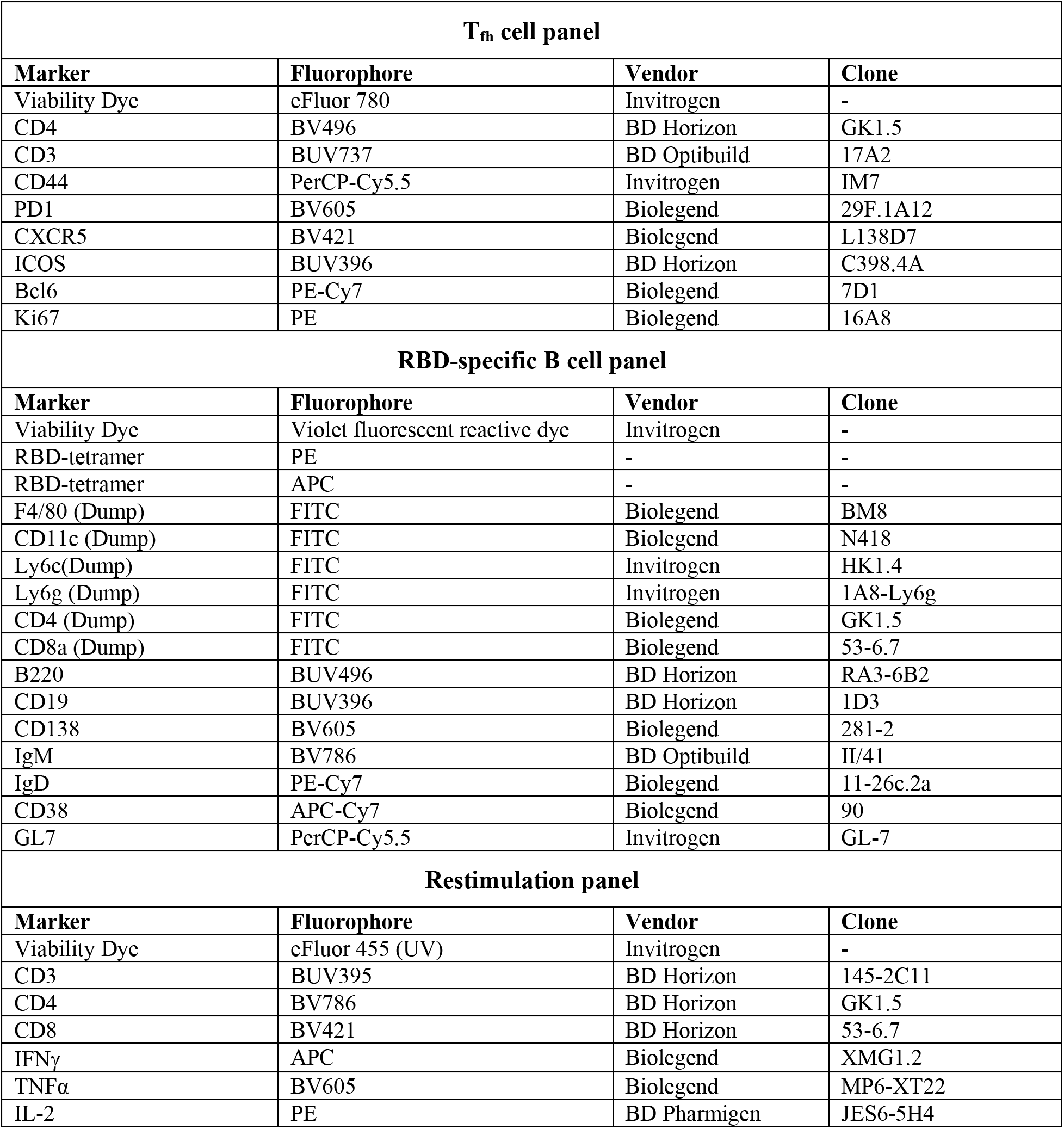
Probes and markers used to characterize cell populations using flow cytometry.

